# New approaches to detecting and characterizing introgression in large species trees

**DOI:** 10.64898/2026.05.30.728990

**Authors:** Sarthak R. Mishra, Laia Pomar-Pallarès, Robert Lanfear, Matthew W. Hahn

## Abstract

Many current phylogeny-based methods to detect introgression use samples of species-quartets to detect asymmetries in gene tree frequencies. While this has proven to be an accurate and robust approach, applying it to larger species trees often means having to test dozens to hundreds of quartets across a tree. Furthermore, any single introgression event can have effects on multiple quartets—with no principled way to determine the number of unique events from a set of quartets—and the direction of introgression cannot always be determined from quartet comparisons alone. Here, we present a new approach to detecting introgression using the frequency with which more distantly related clades are attached to one another among a set of gene trees. Testing for introgression between pairs of branches is straightforward using these discordant attachment frequencies. We further show that the direction of introgression can be inferred between any pair of branches separated by at least two internal branches of the species tree, and that theoretical expectations of gene tree frequencies under introgression can be used to accurately determine the number of independent times genes have been exchanged. Application of these methods to data from cichlids and *Drosophila* demonstrate the power of the new approaches.

The DAFT software package is available from: https://github.com/smishra677/DAFT/

## Introduction

Introgression between species is now recognized as a common, important process across the eukaryotic tree of life (Mallet et al. 2016; Taylor and Larson 2019; Aguillon et al. 2022). Much of the increasing appreciation for the ubiquity of introgression is due to the development of statistical methods that take advantage of genome-scale data (reviewed in Hibbins and Hahn 2022; Huang et al. 2025). The combination of large datasets and powerful methods has made it possible to detect introgression affecting both large and small genomic segments, including adaptively introgressed segments (Racimo et al. 2015).

However, the most robust approaches to detecting introgression from phylogenetic data (i.e. where one or a small number of sequences are collected from each of multiple species) require that we use only subsets of the species at a time. In particular, a suite of related methods are based on measuring the asymmetry in frequency of two minor tree topologies (the ones that do not match the species tree), either among reconstructed gene trees (Huson et al. 2005) or among patterns at individual nucleotide sites (Green et al. 2010; Durand et al. 2011). The wide applicability of these methods (and others, e.g. Edelman et al. 2019) comes in part from the ease with which they can be applied to any dataset (using either gene trees or nucleotide site patterns) that can be used to produce unrooted quartets or rooted triplets of species, as there are only three possible tree topologies in these cases. Nevertheless, the fact that only quartets are sampled is also a problem, as there are many quartets that could be sampled from a large species tree, and even one introgression event can affect multiple of these quartets. As a result, application of quartet-based methods to large species trees often leads to complicated results in which it is hard to infer an accurate number and timing of introgression events (e.g. Suvorov et al. 2022). While attempts have been made both to integrate many quartet-based tests into a minimum set of events (e.g. Elworth et al. 2018; Malinsky et al. 2018) or to have model-based inferences from many sets of quartets (e.g. Solís-Lemus and Ané 2016; Rhodes et al. 2021), both approaches have limitations as to the types of events they can detect.

Furthermore, quartets are fundamentally limited in the types of inferences that can be made from them. For instance, many researchers would like to know the direction of gene flow—i.e. the precise donor and recipient branches—not just the species or lineages involved in an introgression event. Such inferences cannot be made from just the topologies of a quartet, though they can sometimes be made either using branch lengths (Yu et al. 2014; Hahn and Hibbins 2019; Forsythe et al. 2020) or from a larger collection of quartets with specific arrangements of gene flow (Pease and Hahn 2015; Solís-Lemus and Ané 2016; Leppälä et al. 2024). In general, there is still a need for methods that can make accurate inferences about introgression from large species trees (and large sets of gene trees), without turning such datasets into a collection of quartets.

Here, we present a new framework that can help to achieve these goals, comprising two new statistical methods and a brief theoretical examination of introgression that helps in the interpretation of these methods (and other methods). The key insight of our approach is that full gene trees (i.e. from all species in an analysis) contain important information about who is exchanging genes and the direction of this exchange, information that is not contained within quartets. One can readily extract this information from gene trees using what we call “discordant attachments”—branches that are sister in a gene tree, but that are not sister in the species tree. A test on discordant attachment frequencies provides a statistical way to determine the branches affected by introgression. Using these discordant attachments, we also present a new method for determining the direction of introgression. Our method can determine the direction of allele exchange in introgression events involving any two branches that are separated by more than a single internal branch, including in cases of bidirectional introgression. Finally, introgression on a single branch can have radiating effects across a species tree, a problem that plagues quartet-based methods. Therefore, we present an explicit coalescent model of introgression, one that helps to describe how these effects spread, and how to distinguish the branch on which gene flow occurs from those that are affected secondarily. We provide a new software package, DAFT (Discordant Attachment Frequency Tool), that implements these new approaches. Using extensive simulations, we test the accuracy of these approaches, also applying them to datasets from cichlids (Gante et al. 2016) and *Drosophila* (Suvorov et al. 2022) in order to clarify the history of introgression among a larger number of species.

## Methods

### Discordant attachment frequency test

Our method requires as input a collection of reconstructed gene trees and a rooted species tree (no branch lengths are required). All trees must be bifurcating—polytomies are ignored. Errors in tree reconstruction could possibly result in incorrect inferences, but for now we assume that both gene trees and the species tree have been constructed without error. We examine cases with error in the Results.

Given these data, DAFT counts the number of times any two clades or branches that exist in the species tree are seen as sister (“attached”) in a gene tree. In general, our test is based on the idea that introgression will increase attachments between non-sister branches (e.g. Figure 1), over and above the effects of incomplete lineage sorting (ILS) alone. In the example gene tree shown in Figure 1A, the tip branch leading to species *F* is sister to the branch leading to species *A*, which is a discordant attachment since this relationship is discordant with the species tree.

**Fig 1.**
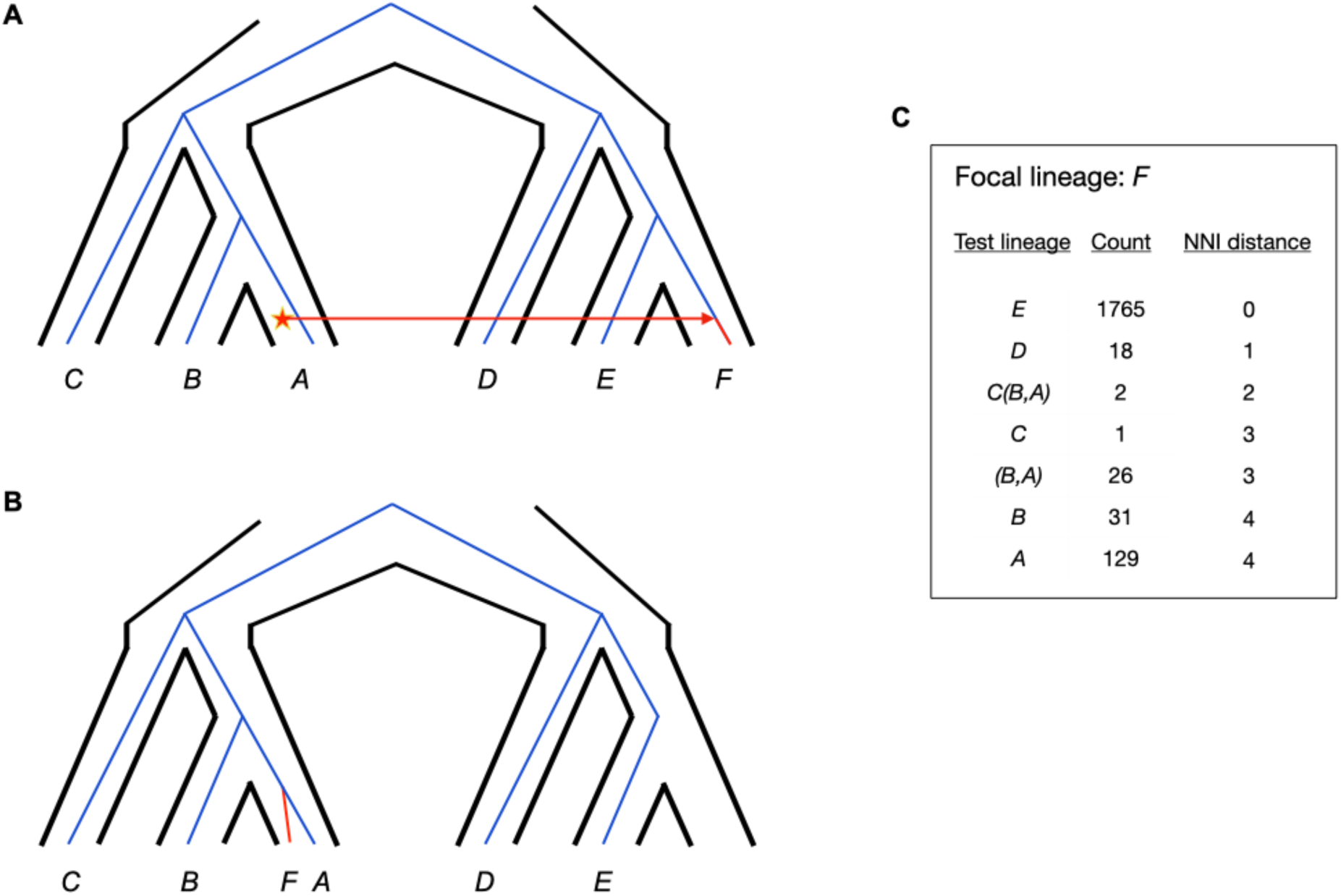
Overview of the effects of introgression on discordant attachments. **A.** A species tree (black outline) with an example gene tree (blue lines) within it. A single introgression event occurs, with donor species *A* and recipient species *F*. **B.** Introgression can lead to discordant attachments, as shown here with an (*A,F*) attachment. The branch leading to species *F* is highlighted in red for clarity. **C.** Example of the type of output generated by DAFT for a simulated dataset with 2000 gene trees. For focal species *F*, its attachments in sampled gene trees to all other branches are counted. Note that only attachments between branches that exist in the species tree are reported by DAFT, which is why the reported attachment counts do not necessarily sum to 2000. Along with a count of the number of times each attachment is observed, the NNI distance between each branch and species *F* is shown. Concordant attachments have NNI distance 0 (i.e. the attachment exists in the species tree), while discordant attachments have larger NNI distances.

There is only one discordant attachment in this gene tree, but multiple such attachments can be counted per tree. Note, however, that even though the clade consisting of (*A,F*) is sister to species *B*, this second attachment is not counted because the (*A,F*) clade does not exist in the species tree (whereas the single-taxon clades *A* and *F* do exist). Discordant attachment counts can be made via a post-order traversal of each gene tree (and in parallel across gene trees), making the calculations relatively fast.

DAFT organizes the data on discordant attachments by individual branch of the species tree. That is, for each branch in the species tree a table of attachments is output (there are 2*n*-2 such branches in a rooted species tree with *n* tips). Figure 1C shows this table for the tip branch leading to species *F*. Among the attachments to a specific focal branch, the data in the table are ordered by the nearest-neighbor interchange (NNI) distance (Robinson 1971; DasGupta et al. 2000) between the focal branch and every other branch in the species tree. The NNI distance between two branches represents the number of NNI rearrangements that must be carried out on the species tree to make them sister to one another (Figure 2); equally, this distance represents the number of rearrangements that must be carried out to move them back to their position in the species tree if they are attached discordantly. Branches with an NNI distance of 0 are therefore concordant with the species tree. The NNI distance is highly similar to the number of internal branches separating two lineages—and is correlated with many other distance measures—but we use it here because it is also useful in inferring the direction of introgression (see next section).

**Fig 2.**
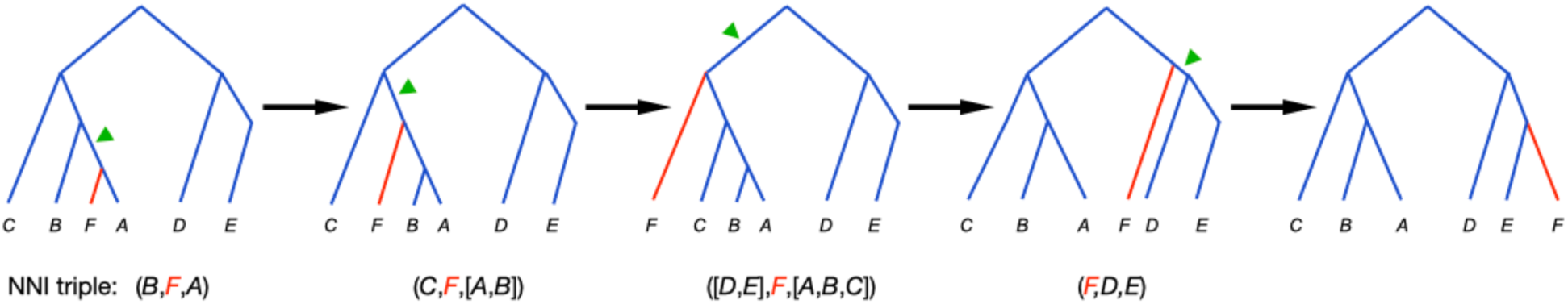
Overview of the djiNNI algorithm. Starting on the left with the discordant gene tree from Figure 1B, the series of NNI rearrangements shown can transform the gene tree into the species tree. A minimum of four NNI moves are required in this case, and each NNI is a rearrangement around an internal branch (highlighted by green arrows). For each internal branch used in an NNI move, we also record the three branches neighboring it (the “triple”), shown below each tree here. After the reconciliation is complete, djiNNI reports the branch that occurred in the most triples—in this case, it is branch *F* (highlighted in red). In this gene tree, therefore, branch *F* is determined to be the one that moved; i.e. it was likely the recipient of gene flow.

Regardless of the exact distance measure used, the organizing principle behind using such a measure is that it is negatively correlated with the number of discordant attachments between two branches expected under ILS alone. For instance, two branches 1 NNI move apart are generally expected to appear together more often than two branches that are 2 NNIs apart.

However, this expectation is true only under certain conditions. First, because NNI distance does not take branch lengths into account, it is not a quantitative predictor of the exact amount of discordance due to ILS. This means that predictions about the relative amount of ILS can only be made along the same path of a species tree. In other words, a pair of branches 2 NNIs apart will have fewer discordant attachments than a pair 1 NNI apart *only if* the internal branches affected in both cases lie along the same path to the common ancestor of the lineage in common between the pairs (Figure S1). Otherwise, we cannot make predictions about relative attachment frequencies.

A second caveat is that internal branches present in the species tree only exist in a gene tree when the descendant lineages have coalesced. If the internal branch of the species tree is long enough this is nearly guaranteed to occur, and expectations about attachment frequency are predicted solely by NNI distance. However, on short internal branches coalescence may not occur, and therefore NNI distance may not predict attachment frequencies for such branches accurately. Consider the internal branch subtending species *A* and *B* in Figure 1. This branch is 1 NNI away from species *D*, and therefore if it is long enough the pair (*A,B*) should have the same attachment frequency to species *D* as does the tip branch leading to species *C*, which also has an NNI distance of 1 from *D*. If their shared internal branch is short, however, species *A* and *B* may not coalesce in any particular gene tree, and there will be a relative deficit of (*A,B*) pairs, leading to fewer attachments of this pair with *D*. In general, when there are very short internal branches the relationship between NNI distance and discordant attachments can be complicated (Than and Rosenberg 2013). Our statistical tests can be corrected in such cases by normalizing by the number of times a branch is seen among all gene trees.

Taken all together, the above principles suggest several straightforward ways to test for introgression. A comparison of sister branches is the simplest test that can be made—for example, Figure 1C shows that the count of (*A*,*F*) attachments (*n*=129) is substantially larger than the count of (*B*,*F*) attachments (*n*=31). These numbers can be used in a statistical test of a pure ILS model, similar to the Δ test of Huson et al. (2005). We have found that these are somewhat noisy tests, with relatively frequent false positives and false negatives, though still informative (see Results). As an alternative, we introduce a test that compares attachment counts between branches that have different NNI distances to a third species. As an example, the pair (*A*,*F*) has an NNI distance of 4, while the pair (*C*,*F*) has an NNI distance of 3 (Figure 1).

Therefore, the strong prediction is that species *C* will attach to species *F* more often than *A* does. Figure 1 shows that this is clearly not the case in our illustrative example: the pair (*C*,*F*) is observed in 1 attachment, while the pair (*A*,*F*) is observed in 129 attachments. As *A* and *B* are referred to as sister branches, we refer to *C* as an uncle branch of *A*; we therefore call these “avuncular” comparisons.

In general, for tests involving focal branch *Z* (branch *F* in the above example), with branch *X* as the test branch (*A* in the example) and branch *Y* as the uncle branch (*C* in the example), we refer to the count of (*X*,*Z*) attachments as *n_test_* and the count of (*Y*,*Z*) attachments as *n_uncle_*. Then, a *z*-score for the avuncular test can be calculated as:

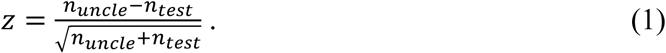

This DAFT uncle test statistic should follow a standardized normal distribution (Pocock 2006). Table S1 shows the results of avuncular tests for the focal branch leading to species *F*.

One advantage of these tests is that we can easily see when the test species has a higher attachment count: the *z*-score is arranged so that the statistic is negative when there is evidence for introgression and positive otherwise. As the null hypothesis under a model with no introgression is that the *z*-score should be positive (i.e. that the lineages separated by fewer internal branches have more attachments), we can also consider carrying out avuncular comparisons as one-sided tests. Although avuncular tests may have lower power than sister-branch tests in certain circumstances, they can have higher power in other circumstances; they also have fewer false positives (see Results). It should be noted that not every test branch allows for an avuncular comparison. In particular, when only one internal branch separates the focal and test branches, no “uncle” branch exists. In these cases, a sister-branch comparison is the only possibility. DAFT carries out all available avuncular and sister comparisons automatically.

DAFT must correct for non-coalescence in comparisons involving short internal branches (as mentioned earlier). In these cases, DAFT can be run by scaling down discordant attachment counts to the lower total number of branches observed in the dataset. For example, above we considered a test comparing (*C,D*) and (*AB,D*) discordant attachment frequencies (cf. Figure 1). If the branch subtending (*A,B*) is short, there will be few gene trees with these together, and even fewer (*AB,D*) discordant attachments. To correct for this, DAFT counts the total number of trees with (*A,B*) pairs, regardless of who this pair is further attached to. If this number is lower than the total number of trees containing branch *C*, DAFT will reduce the number of discordant attachments involving *C*, proportional to the fraction gene trees with (*A,B*) pairs. The DAFT test runs rapidly in correction mode on any dataset.

Two additional aspects of these tests should be mentioned. First, it is often the case in genome-scale datasets that sequences from one or more species will be missing from a gene tree. While randomly distributed missing data should not cause a problem, if one taxon is missing a disproportionate amount of the time this could lead to false signals of introgression. For example, if species *B* in Figure 1 was missing from a large number of gene trees, this might result in an excess of (*A,C*) discordant attachments. Although our method for correcting for non-coalescence should also correct for these missing taxa, users can also re-run DAFT requiring that the specific set of taxa used in the significant avuncular (or sister) test be present in all gene trees to ensure that inferences of introgression are not artefacts in such cases. Second, there is nothing in the results of the avuncular test that provides information on the direction of introgression. Although the results in Figure 1C are shown with respect to branch *F* as the focal branch, DAFT outputs the results of tests from every branch as the focal branch (including internal branches). When the tip branch leading to species *A* is used as the focal one, the uncorrected count of (*A,F*) attachments will be exactly the same, though the *z*-score may be different because species *F* has a different avuncular comparison (species *D* in this case). In order to infer the direction of introgression, in the next section we introduce a second new approach.

### Determining the direction of introgression

It is widely appreciated that the direction of introgression cannot be inferred from *D* or Δ tests, as the same tree asymmetries in a quartet can be generated by both directions of introgression (e.g. Martin et al. 2015). However, in trees with more tips it becomes readily apparent which is the donor and which is the recipient branch. Consider the example in Figure 1: the observed discordant attachment between branches *A* and *F* would not be informative about donor and recipient relationships in a quartet-based test, regardless of which other lineages were used. But in a gene tree with more tips, one can see that the sample from species *F* has been “moved” to the wrong place in the tree, because it is now nested within a clade that also contains species *B* and *C*. This pattern strongly implies that species *A* is the donor, as species *F* has inherited genetic material from a lineage on the other side of the tree.

The above intuition about the effects of the direction of introgression can be turned into an algorithm for determining this direction. Our algorithm, djiNNI (“jē-nē”), which is contained within the DAFT software package, requires the same input data as the test for introgression—gene trees and a species tree—but also requires that the user specify which pair(s) of branches to determine the direction between. We generally presume that users will first run the DAFT test for introgression among all branches (or some other test for introgression), examine those results carefully, and then choose which pairs to run djiNNI on. We advise against attempting to run djiNNI on all possible pairs of branches, to limit the risk of false positive inferences from the very large number of possible tests.

We explain djiNNI for a single focal pair of branches first. In this example we use the terminal branches leading to *A* and *F* (Figures 1,2); however, the method applies equally well to any pair of branches separated by at least two internal branches on the species tree, and need not be limited to terminal branches. The software proceeds initially by scanning all gene trees to find those in which the focal pair are attached—e.g., (*A,F*) trees. For any particular gene tree with this discordant attachment, djiNNI carries out gene tree-species tree reconciliation using only NNI events. This reconciliation transforms the gene tree topology into the species tree topology using a series of NNI moves (Figure 2); such a transformation can be carried out for any two trees with the same set of tips (Robinson 1971; DasGupta et al. 2000). Here, djiNNI takes advantage of optimized software for reconciliation we previously developed that included duplication, loss, and NNI (Mishra et al. 2024).

Any NNI event applied to an internal branch of a rooted gene tree is associated with three flanking branches (a “triple”); these are the branches that are rearranged. We store the clades in each triple for each NNI event needed to transform a specific gene tree into the species tree (Figure 2). Importantly, keeping track of the triples helps to track which branch was moved by introgression: we can see in the example in Figure 2 that the tip branch leading to species *F* is in more triples than the tip branch leading to species *A*. Its presence in more triples implies that it was the branch that received the introgressed DNA, and therefore was the branch that moved farther from its location in the species tree in this particular gene tree. For each gene tree, djiNNI records the number of triples the two focal branches appear in, noting in the end which branch appears in more (there can also be ties, in which case one lineage is picked at random).

Across many gene trees, the expectation is that the branch receiving gene flow will be the one that appears in more triples in more gene trees (for each gene tree the branch that appears in more triples is counted once). The output of djiNNI includes the number of trees in which each of the pair of focal branches was found in more triples, and therefore the branch inferred to be the recipient of gene flow. Of course, introgression is not the only process that can result in branches appearing in the “wrong” place on a gene tree. ILS will cause many rearrangements, including in gene trees that are also affected by introgression, possibly in non-focal branches. In cases involving ILS alone, the expectation is that any two branches appearing in a discordant attachment will appear in approximately equal numbers of triples. Nevertheless, djiNNI does not determine directionality for all pairs of branches, only significant pairs from the DAFT test, so no statistical test is carried out. The branch with the higher count in the focal pair is simply designated as the one that received genetic material (or *more* genetic material; see below), and this branch is not required or expected to necessarily appear in significantly more triples than its pair. The lineage with the higher count is denoted the “major” recipient branch and the one with the lower count as the “minor” branch in the pair.

One consequence of the way inferences are made in djiNNI is that the direction of introgression cannot be determined for pairs of branches with only one internal branch separating them. This occurs because there is only one NNI possible, and therefore only one triple—neither branch can appear more often than the other. Put another way: we can distinguish donor and recipient branch only when they are separated by two or more internal branches (cf. Solís-Lemus and Ané 2016). This rule is also consistent with previous work on the *D* and Δ tests.

It is also possible that introgression is bidirectional, such that each branch in a pair acts as both donor and recipient. For example, we could imagine that in addition to migrants originating from tip branch *A* into branch *F*, migrants also went the other way. Such gene flow would also lead to (*A,F*) discordant attachments, but would act to “move” samples from species *A* into the clade containing species *D* and *E*. Bidirectional introgression does not have to be symmetrical, and therefore we do not necessarily expect the same number of branches moving in the two directions. Therefore, to detect such cases, djiNNI carries out a second analysis, but only for pairs in which the minor branch had more triples than the major branch in at least 10 gene trees in the initial comparison to ensure statistical power. In these pairs, djiNNI asks if the minor branch (*A* in this example) has moved more than expected toward the major branch (*F* here), by carrying out the same analysis for its sister branch (the tip leading to species *B*) attached to the major partner. This sister branch represents the number of moves only due to ILS. Table S2 shows the output from djiNNI for a bidirectional introgression scenario, complete with the *z*-test comparing the relevant counts. If the *z*-test is significant, djiNNI concludes that introgression has been bidirectional (the sign of the *z*-statistic no longer has a meaning). Note that there are some cases in which the bidirectional test cannot be carried out because of the lack of an appropriate sister branch (see Results).

In the end, djiNNI outputs a phylogenetic network (in Extended Newick format) summarizing all of the input pairs and the inferred direction of introgression between them, if this can be determined. However, we have found that single introgression events—even events on tip branches—can result in multiple significant pairs, an effect that can be seen in previous empirical studies using the *D* test (e.g. Malinsky et al. 2018). In order to help to interpret such scenarios, in the next section we outline a coalescent model for introgression.

### A coalescent model for the effects of introgression

Our main goal in the following model is to be explicit about the identity and frequency of gene trees that introgression generates. If we can do this, then we can more precisely identify the correct donor and recipient branches. As our model, we use the parent-tree approach to the multispecies network coalescent introduced by Meng and Kubatko (2009). Although this model has been used to detect and characterize introgression in many previous papers (e.g. Gerard et al. 2011; Liu et al. 2014; Blischak et al. 2018; Hibbins and Hahn 2019; Kubatko and Chifman 2019; Hibbins et al. 2020; Hibbins and Hahn 2021; Allman et al. 2024; Hibbins and Hahn 2024; Dinh and Baños 2025; Kong et al. 2025), this has only been done in the context of a rooted three-species tree (i.e. unrooted quartets). Here, we simply extend the model to a rooted four-species tree, an extension that reveals several interesting patterns that are not evident with fewer taxa.

Assume the species tree shown in Figure 3A, with the time of speciation events (*t*_1_, *t*_2_, *t*_3_) given in coalescent units. We imagine a single instantaneous introgression event from species *A* to species *F* occurring at time *t*_m_. At this point in time, a fraction, γ, of sampled lineages in species *F* have an introgressed history—i.e. they inherit introgressed genetic material from *A*. Under the model used here, non-introgressed lineages (the remaining 1-γ) follow the speciation history (parent tree 1; Figure 3A), while introgressed lineages follow the introgression history (parent tree 2; Figure 3B). Within either history, gene trees are generated according to the multispecies coalescent model (Hudson 1983; Tajima 1983; Pamilo and Nei 1988), with all shared speciation events occurring at the same time in both histories. The frequency of all 15 possible gene tree topologies can be calculated easily for either history (Rosenberg 2002); the individual probabilities are shown in Table S3.

**Fig 3.**
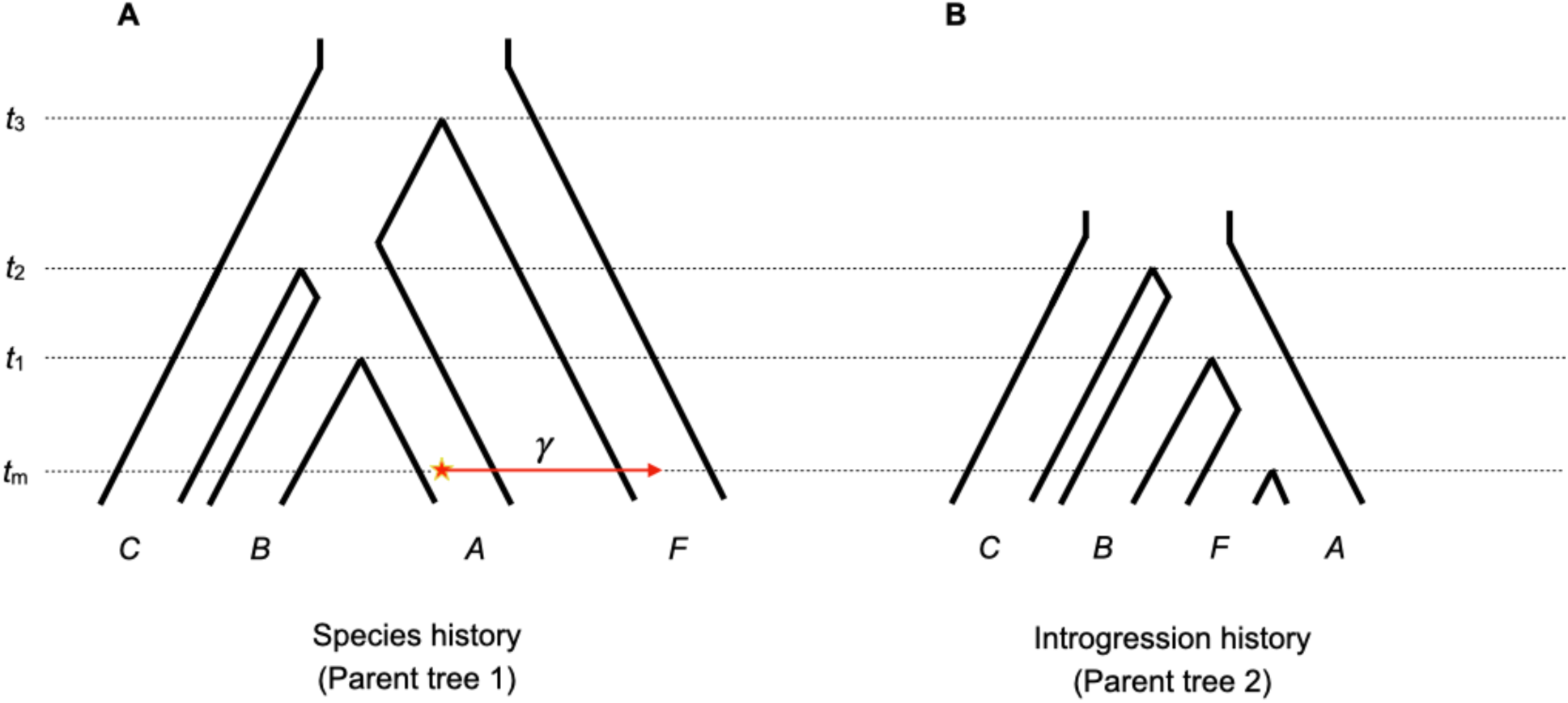
Coalescent model of introgression. The parent-tree approach attempts to model the gene trees resulting from an introgression event using a series of separate parent trees, within which only incomplete lineage sorting can act. **A.** The history of speciation among taxa is modeled with parent tree 1. For the introgression event shown here (red arrow, with the star denoting the donor population), the probability of a gene tree branch in species *F* introgressing is γ. Therefore, parent tree 1 represents the fraction, 1-γ, of non-introgressing gene trees. Non-introgressed gene trees are drawn from the multispecies coalescent using this history. **B.** The introgression event in panel A induces a single additional parent tree, shown here as parent tree 2. The fraction, γ, of gene trees experiencing introgression are all drawn from this parent tree under the multispecies coalescent.

The main effects of introgression on gene tree frequencies using this model are presented in the Results. However, an example may help to make our results more understandable. Suppose we want to know the expected fraction of (*A,F*) gene trees under the scenario shown in Figure 3, with *t*_1_=1, *t*_2_=2, *t*_3_=3, *t*_m_=0.4, and γ=0.1. We first find the expected probability of such trees arising under parent tree 1. There are three different topologies that have *A* and *F* sister to one another, so we need to sum their expected frequencies:

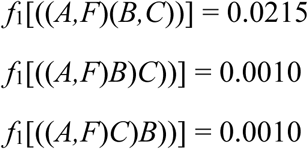

We will refer to the sum from parent tree 1 as *f*_1_(*A,F*) (= 0.0235 here).

Similarly, there are three different topologies that have *A* and *F* sister to one another under parent tree 2 (the introgression history). Note, however, that these are different probabilities because some of the relevant relationships and speciation times have changed:

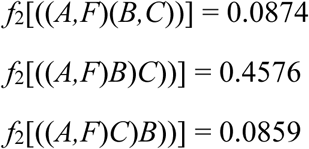

We will refer to the sum from parent tree 2 as *f*_2_(*A,F*) (= 0.6310 here).

Then, to find the total expected frequency of trees uniting *A* and *F*, we sum the contributions from parent tree 1 and parent tree 2, weighted appropriately by γ:

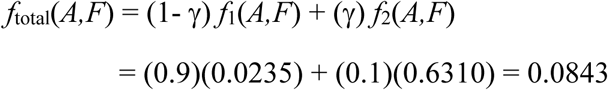

As we can see, there are some (*A,F*) trees expected even without gene flow (i.e. through the speciation history), but a much larger proportion of such trees generated under the introgression history. For comparison under the same scenario, the expected frequency of (*B,F*) trees is:

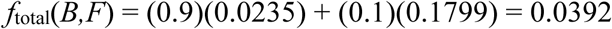

Here, the contribution of (*B,F*) trees from parent tree 1 is the same as (*A,F*) trees, because they are equally affected by ILS in the speciation history. The contribution of (*B,F*) by parent tree 2 (i.e. *f*_2_(*B,F*)) is lower than (*A,F*) trees generated by this history, but is non-zero, which will become important below (see results on the “tail” of introgression). The expected frequency of any particular discordant attachment can be calculated in the same manner—with some modifications for introgression from and to internal branches—given any values of the relevant speciation and introgression parameters. These expected frequencies can then be used to help us to interpret patterns of genealogical heterogeneity in real data.

### Simulations

To test the power and accuracy of the methods introduced here, we carried out simulations using msprime (Baumdicker et al. 2022). We used three different species tree topologies: a 6-species symmetric tree (Figure S2A), a 6-species asymmetric tree (Figure S2B), and a 23-species tree (Figure S2C; based on the tree of non-strepsirrhine primates from Vanderpool et al. 2020). The branch lengths of all topologies are specified in generations, such that varying the (shared) population sizes will change the amount of ILS. For each topology a single haploid sample was taken from each species, and the true tree topologies for each of 2000 gene trees per parameter combination were used unless specified otherwise.

Simulations covered a wide range of conditions. Replicates with different numbers of introgression events (=0, 1, 2), varying levels of introgression (γ=0.02, 0.10), and either high, medium, or low ILS (*N*_e_=10000, 1000, and 100, respectively) were used. In order to add gene tree error to simulations, we added a Poisson number of NNI rearrangements to each tree, with a mean of 0.25 rearrangements for 6-species trees (i.e. on average 25% of trees have a single rearrangement) and 2.5 rearrangements per gene tree for the 23-species tree. The number of rearrangements was drawn separately for each gene tree, and NNI events were applied sequentially so that the same branch could be affected multiple times. Simulations with no introgression at all are used to characterize false positive rates for the DAFT test; simulations with introgression are used to estimate the true positive rate (i.e. accuracy) for the DAFT test and djiNNI.

Simulations with one introgression event varied according to: whether events were tip-to-tip branch, tip-to-internal branch, internal-to-tip, or internal-to-internal (at two different depths of the tree); the NNI distance between donor and recipient (=2, 4, 6); and the timing of introgression on the donor branch relative to the nearest speciation event above it (i.e. closer to the root; *t*_m_/*t*_s_=0.1, 0.5, 0.9). Simulations with two introgression events can have these events arranged in multiple different ways. The three general arrangements used here are: non-overlapping (events do not share a node or edge, and neither is ancestral to the other), overlapping (the two events share edges, or occur on edges lying on the same path to the root), and bidirectional (the two events occur between the same two branches of the species tree, but in different directions). Details on each of the 51 different simulation conditions are given in Table S4.

## Results

### Accuracy of the DAFT test with no introgression

We begin by characterizing the rate of false positives for the DAFT test in simulations without gene flow. These simulations were carried out on three different species trees, with three levels of ILS, with and without gene tree error; these conditions correspond to simulations A-L in Table S4. 10 datasets with 2000 gene trees each were simulated for each condition.

For the symmetric and asymmetric species trees with low and medium ILS, we did not detect a single avuncular test that was significantly negative (*z* < -1.96); this corresponds to a false positive rate (FPR) of 0% (Table S4). In other words, the discordant attachments between branches with higher NNI distance was never significantly greater than between branches with a lower distance. Out of 330 sister-branch tests conducted across these two trees with low and medium ILS, we found 11 significant at *z*>|1.96|. This corresponds to an FPR of 2.7-5% per condition (Table S4). Similar results were found for the 23-species tree with low and medium ILS, with a 0% FPR for the avuncular test and a 0.5% FPR for the sister test.

In the low and medium ILS conditions the shortest internal branches of the species trees are either 11 or 1.1 coalescent units, respectively (Figure S2). In such cases, the statistical correction for non-coalescence in branches sub-tending two or more lineages is not needed (using it did not change any results). In contrast, in the high ILS condition (simulations G, H, and I) the shortest branches can be just 0.11-0.12 coalescent units in all three trees. As a result, the uncorrected FPRs in the high ILS condition across all three species trees were very high (14%-62%; Table S4) for the sister tests. Even in the avuncular tests, where the FPR was lower (1.7-19%; Table S4), on the 23-species tree there were more than a hundred significant tests per simulation. Examination of the results revealed the expected pattern: too few attachments involving clades sub-tended by very short branches, which leads to their sister branches being falsely implicated in introgression. Fortunately, our correction seems to work very well: the corrected FPRs for the same three species trees ranged from 0%-1.7% for the avuncular test and from 2.0%-2.3% for the sister test (Table S4).

In order to examine the effect of gene tree inference error on our false positive rate, we repeated simulations with medium ILS but with error added to each gene tree (simulations J, K, and L; see Methods). We found a small increase in the FPR with this added error (Table S4). In the worst case, for the asymmetric 6-species tree, the false positive rate went from 5% to 13% when one-quarter of the gene trees were inferred in error (simulation K). In the other two cases, the FPR was much smaller (3% for the symmetric 6-species tree, simulation J; 1% for the 23-species simulation, simulation L; Table S4).

### Accuracy of the DAFT test and djiNNI with introgression

For simulations with introgression, we can characterize the true positive rate (TPR) of both the DAFT test (significant or not) and djiNNI (in the correct direction or not). We again vary many factors that might affect the power of these methods, including the level of ILS, the level of introgression, the distance between donor and recipient branches, the timing of introgression relative to the closest speciation event, where on the tree the introgression event occurs, and whether there is gene tree inference error. These conditions correspond to simulations M-TT in Table S4, where all TPRs are also reported.

### Power of sister and avuncular tests

We found generally high statistical power (typically above 80%) in both the sister and avuncular tests. Given that almost all of these simulations had minimal gene flow (γ=0.02), the power of these tests is only expected to go up in scenarios with more introgression. Statistical power is also expected to increase with an increased number of trees sampled (and decrease with fewer trees), but we did not vary this number in our simulations. Here we describe the variables that did or did not have an effect on power, focusing particularly on factors that affect both tests equally; in the next sub-section we describe the conditions under which the avuncular test is more powerful.

Several factors had no or little effect on our ability to detect simulated introgression events in either test. These included gene tree error, which resulted in the same power in the sister test but slightly lower power in the avuncular test (compare simulations CC-FF to M-P). Multiple events occurring in the same dataset had little effect when they were either non-overlapping (simulations MM and NN) or bidirectional (RR, SS, TT). (Note that we considered significant discordant attachments between the introgressing lineages as a true positive, even in bidirectional simulations. Explicit tests for bidirectionality are done using djiNNI below.) The distance between branches involved in introgression, as measured by NNI distance, had little effect on power (compare simulations P, Y, and Z), except in extreme circumstances (see two paragraphs below). While introgressing pairs of branches with smaller distances between them may not have an appropriate uncle lineage with which to carry out the avuncular test at all (simulations Y, GG, II), this depends on the exact shape of the species tree.

Some factors reduced the power of both tests equally. We ran multiple simulations with overlapping introgression events (OO, PP, QQ), some of which had the same branch acting as the donor to two different recipients (simulation PP). In this case, the event that occurred first (i.e. closer to the tips; *Q*→*T*) was detected with high power, but the second event (*Q*→*O*) had lower power. However, this second event was also between lineages that were closer on the tree, which may have contributed to the lower power.

While we generally had very good power when the minimum branch length in a species tree was ∼1 coalescent unit, with very small branch lengths—i.e. large *N*_e_—power was much reduced (simulations Q, R, S, and T). Power is lower because many discordant attachments are seen by chance when ILS is high (for instance, between branches separated by very short distances), drowning out the signal of gene flow. To see what factors could increase the ability to detect introgression with such short branches, we ran additional simulations that either increased the introgression rate (to γ=0.1; simulations Q’, R’, S’, and T’) or increased the NNI distance between the introgressing branches on the 23-species tree (from 4 to 10; simulation T’’).

Increasing the introgression rate generally increased power, for instance going from a TPR of 10% to 60% in simulation T’ (Table S4). Similarly, increasing the NNI distance between the donor and recipient also increased power (simulation T’’), as it reduced the noise due to ILS in a tree with extremely short branches.

Introgression involving internal branches, acting as either donor or recipient, had an effect on power when the introgression event happened shortly after the closest speciation event (looking backwards in time; e.g. simulations KK, LL). To see why this occurs, it is helpful to consider a specific case. In simulated dataset KK, the branch subtending the species pair (*I,C*) is the donor, while the branch subtending species pair (*M,N*) is the recipient (Figure S2C). The signal of introgression is therefore discordant attachments joining (*I,C*) to (*M,N*). While lineages *I* and *C* have had a long time to coalesce before the introgression occurs in this simulation, leading to many gene trees with (*I,C*) branches, introgression happens shortly after speciation of species *M* and *N* (see Figure S3 for a visual explanation of the general issue). As a result, there are very few (*M,N*) pairs that have coalesced, and even fewer discordant attachments between (*I,C*) and (*M,N*). However, there are significant discordant attachments between (*I,C*) and both species *M* and species *N* individually: even though *M* and *N* have not coalesced, both are still affected by introgression (Figure S3). Therefore, it is highly likely that one would still be able to detect this introgression event (see section on “*Inferring the number and timing of introgression events*”).

### The tail of introgression

In several scenarios, the avuncular test has higher power than the sister test (simulations N, AA, BB). This occurs when introgression happens shortly *before* speciation on the donor branch (looking backwards in time), resulting in several patterns that are worth describing in more detail because they occur in many datasets and affect many different introgression statistics.

A helpful place to start is to consider the simple scenario shown in Figure 1A, which corresponds to simulation M (see also Figure 4). The donor in the introgression event is the tip branch leading to species *A*, while the recipient is the tip branch leading to species *F*. Even though introgression is from *A*→*F*, many additional discordant gene trees are observed due to ILS along the donor branch leading to species *A* (Figure 4B-D). Figure 1B shows one outcome of this event, which is a gene tree containing the discordant attachment (*A,F*). However, this is not the only gene tree generated by this introgression: if the tip gene tree branches leading to *A* and *F* do not coalesce before reaching the next speciation event (which occurs at time *t*_1_ in Figure 4A), then branch *F* can coalesce with all three available branches in the next ancestral population (Figures 4B-D). As a result, this single introgression event will also generate discordant attachments (*B,F*) and (*AB,F*).

**Fig 4.**
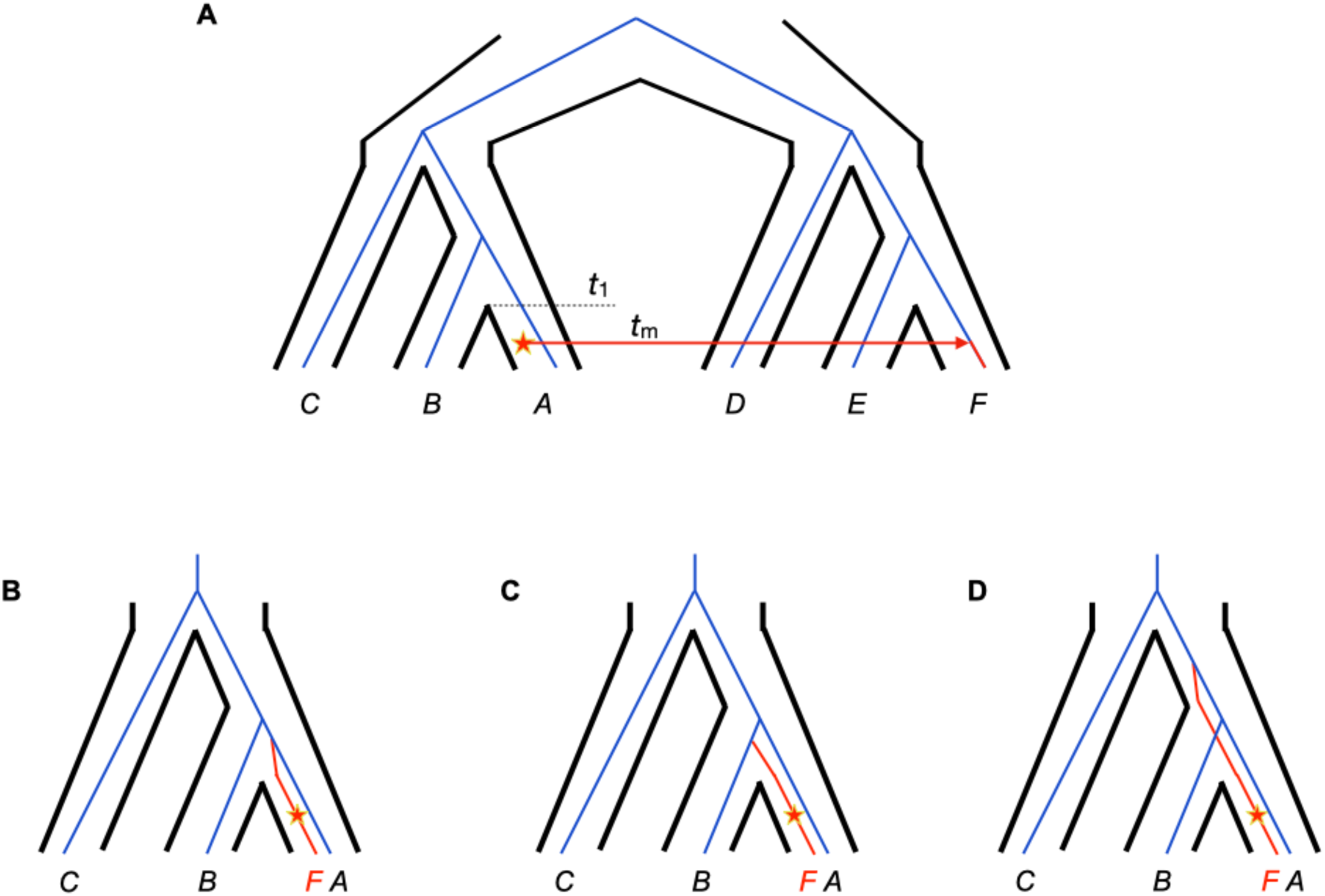
The tail of introgression. **A.** The same species tree and introgression event as in Figure 1 are shown, along with a concordant gene tree. The introgression event occurs at time *t*_m_, while the speciation between species *A* and *B* occurs at time *t*_1_. **B-D.** A close-up of the species tree including only lineages *A, B,* and *C* from panel A. After branch *F* is captured by introgression, it has to coalesce with a donor lineage (looking backward in time). If it coalesces before time *t*_1_, which occurs with probability 1-*e*^-(*t*1-*t*m)^, then it must coalesce with a gene tree branch from species *A* (as shown in Figure 1B). If branch *F* does not coalesce by time *t*_1_, which occurs with probability *e*^-(*t*1-*t*m)^, then it can coalesce with branches available in the common ancestor of species *A* and *B*. The three possibilities shown here are: **B.** Branch *F* coalesces with branch *A*. **C.** Branch *F* coalesces with branch *B*. **D.** Branch *F* coalesces with branch (*A,B*). These three possibilities are equiprobable.

We refer to these further discordant attachments as the “tail” of introgression. This pattern can also be understood from Figure 3, where the parent tree describing the introgression history can generate many additional discordant trees, based solely on the time between migration (*t*_m_) and speciation (*t*_1_). In the Methods, this appeared as the excess (*B,F*) attachments produced by *A*→*F* introgression (i.e. *f*_2_(*B,F*)). If the internal species tree branches were short enough (i.e. *t*_2_ and *t*_3_ close to *t*_m_), *F* could even coalesce with gene tree branches further up the tree, leading to additional discordant attachments.

Why does the tail affect the power of tests of introgression? Simulation N has *A*→*H* introgression in the asymmetric 6-species tree (Figure S2B). Gene flow occurs soon before speciation on the *A* branch, such that ILS on this branch results in approximately equal numbers of (*A,H*) and (*B,H*) discordant attachments. Consequently, sister tests between *A* and *B* are often not significant. However, avuncular tests comparing (*A,H*) with (*C,H*) still retain high power, as the tail in this case does not generate many (*C,H*) attachments. We can also see that moving introgression away from speciation, towards the present (reducing *t*_m_/*t*_s_), increases the power of the sister test (simulation AA), while moving it closer to speciation (increasing *t*_m_/*t*_s_) does not (simulation BB). Similarly, when gene flow occurs in the opposite direction (simulation O; *H*→*A* in the asymmetric 6-species tree) there is no tail of introgression because the tip branch leading to species *H* is relatively long, thus resulting in high power of all tests.

In general, these observations—and the coalescent model of introgression used here (see below)—show how even introgression on tip branches of a species tree can result in signals of discordance among many different branches. As another example, in simulation M, in addition to the expected excess of (*A,F*) discordant attachments, we also observe excess (*B,F*) attachments (Figure 4C; significant in avuncular tests) and (*AB,F*) attachments (Figure 4D; significant in sister tests). Importantly, this tail of introgression does not uniquely affect DAFT: quartet-based tests such as *D* and Δ will also show significant asymmetries in many sampling configurations involving these branches (Figure S4). These discordant gene trees are not false positives per se, but rather genuine signals of the radiating effects of introgression. In the section on “*Inferring the number and timing of introgression events*” we attempt to provide guidelines for identifying the underlying species tree branches involved in introgression in the presence of multiple discordant attachments.

### “Ghosted” introgression

One additional consequence of introgression on genealogical discordance should be mentioned. Although this signal could again be seen as a false positive, and therefore as a negative aspect of multiple different tests for introgression, it is a natural by-product of the introgression process.

As described already, when introgression occurs it moves the recipient gene tree branch toward the donor branch, resulting in a discordant attachment between recipient and donor (or a lineage closely related to the donor). Moreover, this process also removes the recipient branch from its natural position in a gene tree, consequently leaving lineages that have not been affected by introgression behind. As the recipient branch is no longer present in this location, additional discordant attachments occur between the non-affected branches. Figure S5 shows how this could occur between gene tree branches *B* and *C*, resulting in an excess of discordant attachments between them, even though gene flow in this scenario is *F*→*A* (with branch *A* moving toward branch *F*).

This type of effect is well-known when applying quartet-based tests of introgression such as *D* and Δ, at least in cases in which the recipient branch and its non-affected neighbors are sampled (corresponding to branches *A, B,* and *C* in Figure S5). Due to the limited number of sampled tips in quartet-based tests (one of which is usually the outgroup), this effect is only seen when the donor lineage is not sampled; the donor is therefore referred to as a “ghost” lineage (Beerli 2004; Slatkin 2005). The effect of so-called ghost introgression on quartet tests has been explored in depth (e.g. Durand et al. 2011; Ottenburghs 2020; Hibbins and Hahn 2022; Tricou et al. 2022; Tiley et al. 2023; Pang and Zhang 2024). One consequence is a signal of introgression in such tests between lineages that have not themselves been involved in the introgression event.

In the context of a larger species tree, all lineages could have been sampled and we might still see this effect. For example, in simulation O (*H*→*A* in the asymmetric 6-species tree) we find a significant excess of discordant attachments between *B* and *C* in nearly every dataset, because gene tree branch *A* is discordantly attached to branch *H* in every dataset. As the donor lineage is sampled in every gene tree using DAFT, but the recipient has left its close relatives behind, we refer to this as “ghosted” introgression. In fact, such an effect is also seen when applying quartet-based tests to larger species trees (e.g. between branches *E* and *D* in Figure S4), and has little to do with donor lineages being unsampled or extinct: they are simply not included in every quartet. Expectations about the genealogical effects of introgression must take such patterns into account, either in quartet-based approaches or in DAFT. However, note that in the DAFT context these patterns may be directly detectable because the signal of ghosted introgression only appears in gene trees that have an introgressed history, and therefore that also show discordant attachments of the true donor and recipient lineages.

### The accuracy of djiNNI

Our algorithm for determining the direction of introgression, as implemented in djiNNI, was perfect in nearly all tested scenarios (Table S4). While it cannot determine the direction of introgression when donor and recipient are only 1 NNI apart, in almost all other cases it correctly labeled these lineages.

In a few cases in which the signal of introgression was weak (e.g. simulations T, GG, PP), djiNNI was less accurate. This occurred because there were very few discordant attachments due to introgression (relative to those due to ILS), and consequently the signal of the direction of introgression was also weak—there were simply few informative gene trees.

A more interesting case occurred in some of the simulations with bidirectional introgression. In these scenarios, we considered the two directions of introgression as separate events for djiNNI to infer. While one of the directions of gene flow was always inferred with high accuracy, sometimes the other direction was not inferred at all. In these cases, the bidirectional test was not possible. For instance, in simulation SS, which had bidirectional introgression between species *H* and *A* (Figure S6), djiNNI inferred *H*→*A* introgression 100% of the time, but could not infer introgression in the other direction. This occurs because species *H* does not have a sister branch that can independently attach to *A* for use in such comparisons.

While the bidirectional test is therefore not possible in this case, these still contribute to the lower TPR in Table S4. Note, however, that we do detect *B*→*H* movement, as this discordant attachment due to the tail of introgression is informative on its own.

### Inferring the number and timing of introgression events

The above results make it clear that even relatively simple introgression events lead to a multiplicity of discordant gene trees. While many of these gene trees are truly due to introgression, researchers are often interested in the smaller number of biological events underlying these topologies. Characteristics of these events could include the number of events, as well as the identity of donor and recipient species tree branches. In this section, we use our coalescent model to present quantitative and qualitative guidelines for inferring biological scenarios of introgression. Although it is especially difficult to distinguish between events that occur just before or just after speciation, for many other events these guidelines should allow researchers to make clear inferences from data, whether using DAFT or other methods for interrogating gene tree discordance. We organize these guidelines by the types of lineages involved in introgression.

### Recipient is a tip branch

When the recipient of introgression is a tip branch of the species tree, expectations are fairly straightforward. With only a single tip sample, there is only one gene tree branch that can be captured by introgression (i.e. branch *F* in Figure 1A). In terms of the parent-tree model, this scenario requires only a single additional parent tree (Figure 3B), which carries weight γ (see Methods). In terms of Figure 1, branch *F* will be the only recipient branch that appears in discordant attachments, even if there are multiple donors detected (see next sub-section). This makes the recipient branch in such scenarios easy to identify.

We can also use this simple case to demonstrate the quantitative signal of ghosted introgression. We use the introgression scenario shown in Figure S5A (*F*→*A* introgression) and the parent-tree model to calculate the expected frequency of (*B,C*) discordant attachments (e.g. Figure S5B). Figure S5C shows these frequencies, which increase linearly with γ (the probability of introgression). In fact, they increase at nearly the same rate as the frequency of (*A,F*) attachments in this example, since they are a direct and inevitable result of gene trees affected by this introgression event.

### Donor is a tip branch

When the donor of introgression is a tip branch of the species tree (or internal branch, see below), discordant gene trees display the tail of introgression explained earlier. In this case, the probability of non-donor branches displaying excess discordant attachments is determined by *t*_1_-*t*_m_, the time between the next speciation event and introgression (Figure 4A). The greater this time, the less likely a tail will be observed at all. Figure S7A uses the parent-tree model to calculate the expected frequency of discordant attachments under the introgression scenario shown in Figure 4 (*A*→*F* introgression). When *t*_1_-*t*_m_ is large, (*A,F*) is the only discordant attachment found at higher-than-expected frequency (Figure S7B). As this quantity gets smaller, multiple additional discordant attachments become more frequent, until the point at which (*A,F*) and (*B,F*) trees are equiprobable.

Overall, these expectations suggest that introgression from a tip branch should produce discordant gene trees involving this donor branch at higher frequencies than other branches (i.e. (*A,F*) at the highest frequencies). However, as seen in the results from simulation N, as introgression gets close to speciation, one might observe equal (or even more) counts of (*B,F*) or (*AB,F*) attachments stochastically. The timing of introgression and exact donor in these cases will therefore be difficult to identify with certainty.

### Recipient is an internal branch

When the recipient of introgression is an internal branch of the species tree, the parent-tree model is not as easy to specify (Yu et al. 2011; Degnan 2018). This occurs because multiple different gene tree lineages (and gene tree topologies) can be “captured” by the introgression event if this occurs close to speciation (i.e. *t*_m_-*t*_1_ is small; Figure S3). As a result, even a single introgression event must be accommodated by multiple parent trees.

In the Supplementary Note (S1 File) we show how the parent-tree model can be used when the recipient of introgression is an internal branch of the species tree. For an internal branch with any number of descendants, we show how to calculate the number of alternative parent trees that must be added alongside the species tree, a number that can grow quite high when internal branches subtend many tips (approximately as the cube of the number of tips). For internal branches subtending only two tips, six additional parent trees are necessary for accurate expectations: two for non-introgression and four for introgression histories. These parent trees are shown in Figure S8 for *F*→(*A,B*) introgression. The introgression histories (Figure S8D-G) include instances in which gene tree branch *A*, *B*, (*A,B*), or both *A* and *B* are captured by introgression (the last possibility occurs with frequency γ^2^).

Using this model, we can calculate the expected frequencies of discordant attachments, demonstrating that these expectations match simulations very closely (Figure S9). As anticipated, the frequency of discordant attachments involving (*A,B*) goes up as *t*_m_-*t*_1_ gets larger, since more of the gene trees on the recipient branch have coalesced before introgression. Closer to speciation (i.e. *t*_m_-*t*_1_ small) we see discordant attachments involving *A*, *B*, and (*A,B*) almost equally. Importantly, gene tree branches *A* and *B* are always equally represented, which helps to pinpoint the internal branch (as opposed to a tip branch) as the recipient of introgression.

Nevertheless, a short time between speciation and introgression—coupled with possibly small numbers of gene trees showing signals of introgression—may make it difficult to distinguish among multiple possibilities.

### Donor is an internal branch

When the donor of introgression is an internal branch of the species tree, many of the processes explained for other scenarios may all act at once. For instance, in the case of (*A,B*)→*F* introgression (Figure S10A), the tail of introgression can still act (when *t*_2_-*t*_m_ is small) resulting in discordant attachments possibly involving branch *C*, even though the donor is branch (*A,B*). Even if coalescence happens long before the next speciation event (looking backwards in time), it can happen shortly after the previous speciation event (*t*_m_-*t*_1_ small; Figure S10A), which means that *A*, *B*, or (*A,B*) can be involved in discordant attachments with recipient lineage *F* (Figure S10B,C). Again, *A* and *B* are expected to be equally represented among discordant attachments.

A multiplicity of discordant attachments are possible when both the donor and recipient branches are internal. Consider the scenario shown in Figure S3, which has internal species tree branch (*A,B*) as the donor and internal branch (*E,F*) as the recipient. If introgression occurs shortly after speciation on both branches, the donor lineage can be represented by gene tree branches *A, B*, and (*A,B*), while the recipient lineage can be represented by gene tree branches *E, F*, and (*E,F*). Discordant gene trees can then involve attachments between any pair of donor and recipient branches. If introgression on the donor branch occurs soon before the next speciation event, then species *C* gene tree branches may also appear in discordant attachments due to the tail of introgression.

### Analysis of biological data

To demonstrate the utility of DAFT, we use two datasets with available gene trees that have explored histories of introgression. In both cases, previous authors used some combination of quartet-based tests and other analyses in order to propose a set of introgression events. As these authors are organismal experts and are familiar with many aspects of these systems, our goal is not to provide contradictory (and possibly biologically implausible) histories. Instead, we hope that the application of the framework introduced here provides clear hypotheses that can be evaluated in the future.

### Cichlids

Gante et al. (2016) explored the history of speciation and introgression in *Neolamprologus* “Princess” cichlids from Lake Tanganyika. Figure 5A here represents the authors’ inference of the timing and direction of introgression events among five of these cichlid species (two outgroups are not shown). While these authors used gene trees, site patterns, and other information jointly to infer this history, here we used only 4,781 gene trees from this paper as input to DAFT.

**Fig 5.**
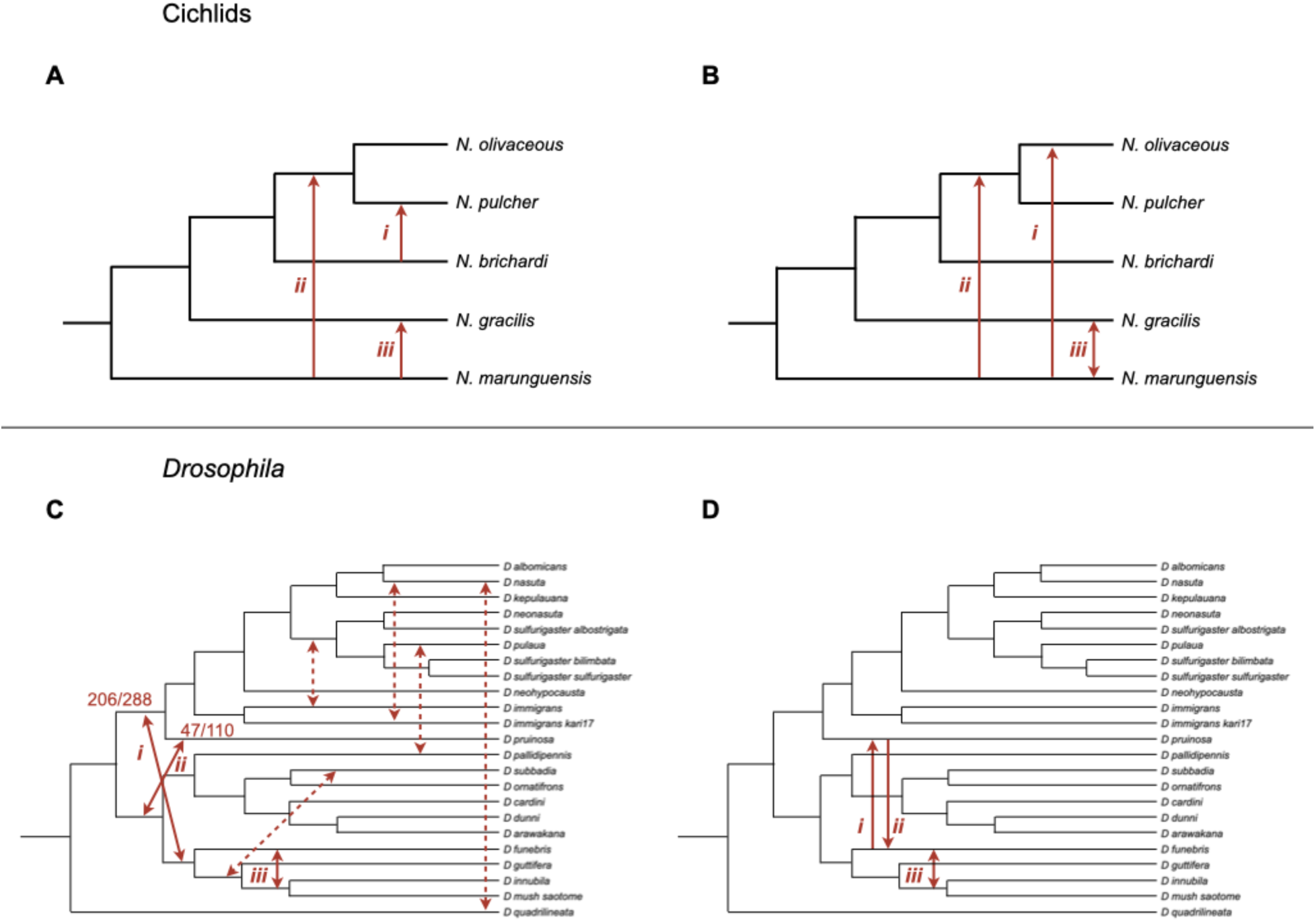
DAFT clarifies previous empirical results. **A.** The introgression history of *Neolamprologus* cichlids originally inferred by Gante et al. (2016). Three introgression events were inferred by these authors using a combination of methods. These events are labeled with lowercase Roman numerals. Note that neither the species tree branch lengths nor the placement of introgression arrows are intended to impart any information about the absolute timing of these events. **B.** The introgression history inferred by DAFT and djiNNI using 4,781 gene trees from Gante et al. (2016). DAFT matches one event exactly (event *ii*), including the direction of gene flow, while for one other it matches the two lineages involved (event *iii*). However, because these lineages are only 1 NNI move apart, djiNNI cannot determine the direction of introgression between them (shown as a double-headed arrow). DAFT also infers a third event that was not found in the original study, between *N. marunguensis* and *N. olivaceous* (event *i*). The introgression between *N. brichardi* and *N. pulcher* in the original study (panel A, event *i*) is likely an effect of ghosted introgression resulting from the event found here. **C.** The introgression history of *Drosophila* “clade 9” originally inferred by Suvorov et al. (2022). Many introgression events were inferred by these authors, using a combination of quartet-based tests. The events shown as dashed lines were supported by only one asymmetric quartet out of all possible quartets. The other events are labeled with lowercase Roman numerals. For two of the other events, the count of significant quartets out of all possible quartets involving those branches are shown (from original). Note that neither the species tree branch lengths nor the placement of introgression arrows are intended to impart any information about the absolute timing of these events. **D.** The introgression history inferred by DAFT and djiNNI using 2,791 gene trees from Suvorov et al. (2022). None of the original events with dashed lines are found. For one event DAFT agrees perfectly with previous results: this is shown as a double-headed arrow because the direction of introgression cannot be determined (event *iii*). DAFT and djiNNI also infer bidirectional introgression between *D. pruinosa* and *D. funebris* (events *i* and *ii*), two events that appear to explain all of the original significant quartets in this part of the tree, due to the tail of introgression.

The DAFT test run on the 4,781 gene trees detected five discordant attachments (all were still significant after correction). Two of these pairs involved branches of the species tree more than 1 NNI apart, and were used as further input to djiNNI. These two events could be clearly inferred to involve unidirectional introgression from *N. maraguensis*, into the tip branch leading to *N. olivaceous* (Figure 5B, event *i*) and into the shared ancestral branch above *N. olivaceous* and *N. pulcher* (Figure 5B, event *ii*). This latter event agrees exactly with one of the events originally proposed in Gante et al. (Figure 5A, event *ii*).

The three additional discordant attachments were 1 NNI apart, and therefore their direction could not be inferred. One of these is shown between *N. maraguensis* and *N. gracilis*; because the direction cannot be determined, this event is shown as a double-headed arrow (Figure 5B, event *iii*). This event was also identified by Gante et al., though they also inferred the direction of gene flow (Figure 5A, event *iii*). However, the other two significant discordant pairs we detected, between *N. pulcher* and *N. brichardi* (also identified by Gante et al.; Figure 5A, event *i*) and between *N. brichardi* and *N. gracilis*, correspond exactly to “ghosted” introgression scenarios involving the detected unidirectional events, and we therefore conclude that they are likely not independent events (and do not include them in Figure 5B).

#### Drosophila

Suvorov et al. (2022) sequenced the genomes of a large number of *Drosophila* species. As the common ancestor of all included species existed quite a long time ago (∼50 million years), their analyses of introgression were conducted on separate sub-clades with more recent ancestors. We confine our analyses to their “clade 9,” which includes 23 species or sub-species, as this clade showed multiple ambiguous introgression events (Figure 5C) that seemed as though they could be clarified with DAFT. We use the same 2,791 gene trees that these authors used to infer introgression using multiple quartet-based tests and the *f*-branch method (Malinsky et al. 2018; Malinsky et al. 2021). Based on the possibility raised by the authors of low accuracy in some gene trees, we only evaluate evidence for the events originally identified in their paper (Figure 5C).

After running DAFT on the 2,791 gene trees, we found multiple pairs of discordant attachments using both the sister and avuncular tests, before and after correction. Multiple events had low confidence in the original study because they were only supported by a single significant quartet (shown as dashed lines in Figure 5C); we found no evidence for excess discordant attachments associated with any of these events. In fact, we found no discordant attachments at all (i.e. no gene trees with these branches sister to one another) between all five of these pairs of clades. These relationships were therefore likely due to events elsewhere on the tree, or were simply false positives.

Among the remaining events, we found both support and clarity. We confirm introgression between *D. funebris* and the ancestor of *D. innubila* and *D. mush saotome* (Figure 5C, event *iii*; Figure 5D, event *iii*). Because these branches are separated by 1 NNI, we are unable to determine the direction of introgression. Suvorov et al. also found many asymmetric quartets closer to the root, among many different tips. For one of the inferred events shown in Figure 5C (event *i*), 206 of 288 possible quartets were significant for introgression between the implicated branches; for the other event (*ii*), 47 out of 110 possible quartets were significant.

Based on the results from DAFT and djiNNI, it appears that all of these significant quartets were generated by bidirectional introgression between *D. funebris* and *D. pruinosa* (Figure 5D, events *i* and *ii*). In both donor lineages, the tail of introgression has resulted in multiple discordant pairs involving closely related branches, resulting in a complex set of asymmetric quartets.

## Discussion

Introgression between species is a complex process. Rather than being an alternative to stochastic processes such as ILS and coalescence, it is instead always accompanied by these processes. Analyses of gene flow are therefore not testing introgression vs. ILS, but rather ILS with introgression vs. ILS by itself (Hibbins and Hahn 2022). With all of these processes acting at once, there can be a large number of gene tree topologies produced by even simple events.

This complexity is further compounded by the stochastic errors and systematic biases introduced when reconstructing gene trees from finite alignments using simplified models of evolution.

As a result of the complexity of introgression, it can be difficult to interpret the genealogical patterns produced by even relatively simple introgression scenarios. While there are a number of powerful, robust tests for the presence of introgression, they often work on subsets of genealogical data, such as quartet trees. Here, we have introduced a new framework for analyses of introgression, one that uses full gene trees to track discordant attachments among possibly introgressing species. This framework, including two separate programs within the software package DAFT and a coalescent model of the effects of introgression on gene tree frequencies, is intended to be used as a whole. However, any part can also be used by itself in conjunction with other tests or analyses, providing assistance in interpreting a wide range of scenarios.

DAFT, and the analysis of discordant attachments in general, has both advantages and disadvantages. The main advantage of the approach introduced here is that it can be much clearer than quartet-based analyses about the introgression events that have occurred. Due to the radiating genealogical effects of gene flow—including the “tail” of introgression and “ghosted” introgression—many relationships can be affected by even a single event (e.g. Figure S4).

Although model-based methods such as SNaQ (Solís-Lemus and Ané 2016) are able to infer more accurate histories of introgression, these histories are often limited to so-called level-1 networks: those that do not have multiple introgression events sharing any branches (Allman et al. 2024). While more complex histories may be inferable in the near future (e.g. Kolbow et al. 2026), many such methods are still limited to input quartet trees. In addition to advantages in inferring an accurate number and timing (and direction) of introgression events, DAFT should also be quite robust to the effects of natural selection, avoiding the possibility of false positives when the assumptions of more complex model-based approaches are violated (cf. Smith and Hahn 2024).

DAFT also has possible disadvantages. The main limitation of DAFT, especially in comparison to quartet-based methods, is that one must infer individual gene trees. Although many quartet-based methods also use gene trees, for more recent divergences the *D* test can be used and only requires nucleotide site patterns (Green et al. 2010). Our simulations including gene tree inference error did not show large effects on the accuracy of DAFT, but more extreme scenarios could of course lead to increased error. On the other hand, the strong agreement found here between published empirical results and DAFT run on the same datasets suggests that levels of gene tree error experienced in many datasets may not be problematic for DAFT inferences. In addition, the reliance on full gene trees also means that our expectations can depend on coalescence between internal branches—i.e. sub-clade attachment frequencies—a problem that does not occur when only quartets of tips are sampled (though it can occur when calculating gene concordance factors; Lanfear and Hahn 2024). While our statistical correction appears to have accommodated this reliance, running *D* or Δ tests alongside DAFT could be a way to verify any signals it detects. On the other hand, multiple recent studies have stressed the fragility of site-based analyses to variation in nucleotide substitution rates (Frankel and Ané 2023; Koppetsch et al. 2024; Pang et al. 2025). Using full gene trees will be much more robust to such variation, except in extreme circumstances (e.g. Felsenstein 1978).

We have also introduced a new way to infer the direction of introgression, applicable any time the branches exchanging genes are more than one NNI rearrangement away from one another. This method, as implemented in the djiNNI algorithm, is fast, accurate, and does not depend on a particular arrangement of introgressing taxa, as have earlier methods based on quartets (Pease and Hahn 2015; Leppälä et al. 2024). In fact, these earlier methods can be seen as ways to have quartets provide similar information on the branches that are being moved by introgression, albeit very indirectly. In addition, djiNNI can infer bidirectional introgression between pairs of branches, something that is not possible using quartet methods. Nevertheless, this inference is not always possible, and djiNNI does rely on the presence of a sister clade not affected by gene flow in order to detect bidirectional introgression.

The idea of using discordant attachments to study introgression could be extended in the future in multiple ways. One relatively straightforward application should be in estimating the rate of introgression, γ. While it may not be possible to estimate this parameter between branches only one NNI apart (Martin et al. 2015), for introgression among more distantly related species the use of discordant attachments—including attachments due to the tail of introgression—should provide sufficient data. Further, in the current version of DAFT we ignore attachments between clades that do not exist in the species trees. This was done to simplify inferences, despite the fact that these attachments may contain information about introgression. In the future we can imagine using such attachments to learn more about gene flow, for instance the order of events occurring on a single branch. Finally, we can imagine multiple ways to automate inferences of the identity of events implied by the output of DAFT, either by using parametric or non-parametric (i.e. machine learning) approaches. Regardless of additional, potential applications, DAFT currently provides a robust and accurate tool for studying introgression between species.

## Acknowledgments

We thank Wayne Maddison for many helpful tips on the development of the DAFT software package and Leonie Moyle for multiple constructive suggestions on the analyses. Megan Smith first explained the problem of non-coalescence in ancestral branches to us. Anton Suvorov and Dan Schrider helpfully guided us through an explanation of their *Drosophila* data.

## Data availability

The DAFT software package and all code for running the simulations used here is available at https://github.com/smishra677/DAFT/.

## Supporting information

**S1 Fig. Circumstances in which NNI distance predicts relative attachment frequency.**

A species tree with five tips is shown (black lines). For three pairs of branches, we also show the NNI distance between them; i.e. the number of rearrangements required to make them sister to one another. For each of the three pairs, the colored arrowheads indicate the internal branches around which the necessary NNI rearrangements must occur. The pair *C-D* is 1 NNI apart and is predicted to show more attachments due to ILS than pair *A-D*, which is 2 NNIs apart. We can make this prediction because both *A* and *C* share a most recent common ancestor with species *D*. In contrast, even though pair *E-D* is 1 NNI apart, we have no basis for expecting more attachments than pair *A-D*, as they do not share a common ancestor.

**S2 Fig. Species trees used in simulations.**

**A.** A 6-species symmetrical tree. Both external and internal nodes are labeled. Branch lengths are given in generations. Here is the tree in Newick format:

(((((A:300.0,B:300.0)D:1200,C:1500)E:1200,F:2700)I:1200,G:3900.0 )J:1200,H:5100)K;

**B.** A 6-species asymmetrical tree. Both external and internal nodes are labeled. Branch lengths are given in generations. Here is the tree in Newick format:

(((A:300.0,B:300.0)D:1200,C:1500)E:1200,((F:300.0,G:300.0)I:1100 ,H:1400)J:1300)K;

**C.** A 23-species tree, based on the tree of non-Strepsirrhine primates inferred in Vanderpool et al. (2020). Both external and internal nodes are labeled. Branch lengths are given in generations. Here is the tree in Newick format:

(A:10282.960000000001,(((W:2999.210000000001,V:2999.210000000001 )N1:2999.19,(B:2999.210000000001,U:2999.210000000001)N2:2999.19) N3:2999.19,((((K:2142.29,L:2142.29)N4:2142.2799999999997,(M:2142 .29,N:2142.29)N5:2142.2799999999997)N6:2142.2799999999997,(J:514 1.48,((E:2570.74,(D:1285.37,H:1285.37)N7:1285.37)N8:1285.37,((C: 1285.37,I:1285.37)N9:1285.37,(G:1285.37,F:1285.37)N10:1285.37)N1 1:1285.37)N12:1285.37)N13:1285.37)N14:1285.37,(T:6426.85,(S:5141 .48,(O:3856.11,((Q:1285.37,R:1285.37)N15:1285.37,P:2570.74)N16:1 285.37)N17:1285.37)N18:1285.37)N19:1285.37)N20:1285.37)N21:1285. 37)N22;

**S3 Fig. Possible gene tree branches captured by introgression into an internal branch.**

**A.** For the species tree shown (black outline), an introgression event occurs with internal branch (*A,B*) as the donor and internal branch (*E,F*) as the recipient. One specific outcome of this introgression on a gene tree—the capture of branch *F* (highlighted in red)—is shown. **B-D.** Three alternative outcomes of the same introgression event, focusing on the clade of the species tree that contains the recipient branch. Introgression could capture: **B.** the tip branch leading to species *E*; **C.** the single internal gene tree branch leading to the pair (*E,F*); or **D.** both tip branch *E* and *F* independently. See the Supplementary Note for the probabilities of events shown in A-D.

Note that the results for simulated dataset KK discussed in the main text occur in a similar configuration (internal-to-internal introgression), but that pair (*I,C*) and (*M,N*) in that simulation are equivalent to (*A,B*) and (*E,F*) here, respectively.

**S4 Fig. The effects of introgression on quartet-based tests.**

We simulated the introgression event shown on the species tree at the top (*A*→*F* introgression; γ=0.1 and *N*_e_=100) and provided the results to the program Dsuite (Malinsky et al. 2021) to calculate *f*-branch statistics. The output shown here is the *f*-branch (*f_b_*) statistic for introgression between pairs of branches (it is not possible to calculate a statistic for cells in grey). Importantly, the single introgression event simulated here, between two tip branches of the species tree, results in five cells with increased values of *f_b_*. These results represent a mix of the true event—in cell (*A,F*)—results due to the tail of introgression—in cells (*B,F*), (*AB,F*), and (*C,F*)—and results due to ghosted introgression—in cell (*D,E*).

**S5 Fig. How gene flow causes ghosted introgression.**

**A.** A species tree with *F*→*A* introgression. This tree topology also acts as parent tree 1 (no introgression) in the coalescent model used here. **B.** Parent tree 2 for this scenario, which acts as the history of introgression. We also show the most probable gene tree inside this parent tree (blue lines), which has both species pair *A* and *F* (the pair directly affected by introgression) and species pair *B* and *C* sister to one another. We refer to the effect on species *B* and *C* as ghosted introgression, since they can also show a significantly increased number of attachments even though both donor and recipient taxa have been sampled. **C.** The increase in the frequency of (*B,C*) attachments with increasing levels of introgression (γ). Results are expectations from the coalescent model, with the marginal effects on the frequency of all three gene trees containing (*B,C*) also shown.

**S6 Fig. Cases in which tests for bidirectional introgression cannot be carried out.**

A species tree with *H*→*A* and *A*→*H* (i.e. bidirectional) introgression.

**S7 Fig. Frequency of discordant attachments due to the tail of introgression**

For the introgression scenario shown in Figure 4 (*A*→*F* introgression), we used the coalescent model to calculate the expected frequency of different discordant attachments. **A.** The expected absolute frequency of different discordant attachments as a function of the introgression time, *t*_m_. The speciation between species *A* and *B* occurs at *t*_1_=20, and is denoted by the dashed blue line. When introgression occurs long before speciation (*t*_m_<< *t*_1_), attachment (*A,F*) is most frequent. As introgression gets closer to the time of speciation, gene tree branch *F* does not coalesce only with *A*, and additional discordant attachments are observed. We refer to these additional signals as the tail of introgression. **B.** Similar to panel A, but the difference between expected attachment frequencies in scenarios with introgression and without introgression are shown. We refer to this difference (introgression – no introgression) as the excess.

**S8 Fig. The parent-tree model when the recipient is an internal branch.**

**A.** The history of speciation and introgression is shown, with branch *F* as the donor and branch (*A,B*) as the recipient. This tree does not act as a parent tree in this scenario. **B-C.** Parent trees representing histories without introgression. As explained in the Supplementary Note, two separate parent trees are needed to account for cases in which branches *A* and *B* either have not (panel B) or have (panel C) coalesced before the time of introgression. Note that in panel C we represent branches *A* and *B* as a single lineage because they are required to have coalesced. **D-G.** Parent trees representing histories with introgression. As explained in the Supplementary Note, there are four parent trees necessary to represent histories with introgression. These parent trees also correspond to the four scenarios shown in S3 Figure A-D, with branches *A* and *B* here corresponding to *F* and *E* there.

**S9 Fig. Frequency of discordant attachments when the recipient is an internal branch.**

For the introgression scenario shown in S8 Figure (*F*→(*A,B*) introgression), we used the parent-tree model to calculate the expected frequency of different discordant attachments (lines). The x-axis represents introgression occurring at increasing times after speciation (*t*_m_-*t*_1_), with the speciation event denoted by the vertical dashed blue line. As this time increases, gene trees in the recipient branch become more likely to have coalesced, resulting in more discordant attachments of the type (*AB,F*). In addition, we simulated gene trees across a range of value of *t*_m_-*t*_1_, recording the frequency of discordant attachments in each. These results are shown as colored dots corresponding to each attachment type. For each value of *t*_m_-*t*_1_, we simulated 100,000 gene trees in msprime.

**S10 Fig. Frequency of discordant attachments when the donor is an internal branch.**

**A.** The history of speciation and introgression is shown, with branch (*A,B*) as the donor and branch *F* as the recipient. **B.** For the scenario shown in panel A, we used the coalescent model to calculate the expected frequency of different discordant attachments. The x-axis represents introgression occurring at increasing times after speciation (*t*_m_-*t*_1_), with the speciation event denoted by the dashed blue line. As this time increases, gene trees in the donor branch become more likely to have coalesced, resulting in more discordant attachments of the type (*AB,F*). **C.** Similar to panel B, but the difference between attachment frequencies in scenarios with introgression and without introgression are shown. We refer to this difference (introgression – no introgression) as the excess.

**S1 Table. Example of attachments and avuncular tests.**

For the event depicted in Figure 1, we show the attachments for focal branch *F* (as in Figure 1C) and the results of avuncular tests. Only two avuncular tests are possible in this tree, comparing either (*A,F*) and (*C,F*) or (*B,F*) and (*C,F*). In other words, branch *C* is the uncle of both branch *A* and branch *B*. The table also shows the number of times the discordant attachment (*C,F*) is observed and the results of the *z*-test described in the Methods.

**S2 Table. Example output from djiNNI.**

The output from djiNNI with bidirectional gene flow between branches *A* and *F* in the 6-species symmetrical tree. After running tests for introgression in DAFT, the gene trees, species tree, and the identity of the five different discordant attachment pairs have been passed to djiNNI (note that four of these pairs are due to the tail of introgression.) In this case, djiNNI correctly identified that there was bidirectional introgression, as well as the correct donor lineage for all pairs.

**S3 Table. Gene tree probabilities.** Probability of each gene tree topology for species tree (((*Z*,*Y*),*X*),*W*) under the multispecies coalescent (from Table V and Equation 2 in Rosenberg [2002]). Both the speciation history (parent tree 1) and the introgression history (parent tree 2) shown in Figure 3 have this same tree shape, though the length of the two internal branches differs between them. For either history, we denote the length (in coalescent units) of the more recent internal branch as *a* and the older internal branch as *b*. The length of *a* is therefore *t*_2_- *t*_1_ in parent tree 1 and *t*_1_- *t*_m_ in parent tree 2, while the length of *b* is *t*_3_- *t*_2_ in parent tree 1 and *t*_2_- *t*_1_ in parent tree 2.

**S4 Table. Simulation descriptions and results.**

The 51 simulation conditions carried out here. For conditions without introgression, the false positive rate (FPR) is reported for both the avuncular and sister tests, with and without correction for non-coalescence in ancestral populations when necessary. For conditions with introgression, the true positive rate (TPR) is reported for both tests and for djiNNI. In cases with multiple introgression events, we report results from djiNNI separately for each event. In general, different colors indicate different parameters or analyses.

