## Supplementary material for "New approaches to detecting and characterizing introgression in large species trees": S1_File.pdf

May 2026

### 1 Background

We would like to compute the expected frequency of an arbitrary gene tree topology, $\tau$, under a specific species tree with introgression, when the recipient is an internal branch. (All calculations here also apply to the frequency of specific concordant or discordant attachments, with no loss of generality. One just has to sum across all gene trees containing the attachment of interest.) Unfortunately, the approach of the parent-tree model used when the recipient is an external branch—simply add a single additional parent tree representing the introgression history (e.g. Figure 3 in the main text)—does not work. Instead, in this situation a single introgression event requires multiple additional parent trees, depending on the number of ancestral lineages of gene trees that can be present in the recipient branch at the time of introgression.

The key distinction is that, in the case of introgression into a tip branch (and when only a single sample from each species is considered), there is only a single gene tree lineage that can be captured by introgression. On the contrary, when introgression is into an internal branch, the gene tree lineages below this branch may not yet have fully coalesced into a single lineage, and introgression can affect (or not) each of these lineages differently. Even if some of the lineages have coalesced, if there are more than two they may have coalesced in different orders, resulting in different sub-topologies. Therefore, we can have several different "lineage configurations."

### 2 Full calculations with two descendants

We demonstrate the calculations needed to obtain expected gene tree frequencies with an example of introgression into an internal branch with two descendant species, which is the simplest case. Consider the species tree $(((A,B),C),F)$ with introgression from $F$ to the internal branch $(A,B)$ at time $t_m$ (Figure S8). We set the probability of a lineage introgressing to be $\gamma$. At time $t_m$, the recipient species tree branch may exhibit different lineage configurations depending on which coalescent events have occurred.

#### 2.1 Lineage configurations

Given that there are two species below the recipient branch, there are only two possible lineage configurations:

$S_1$ : The lineage configuration in the recipient branch at time $t_m$ consists of two lineages, $\{A, B\}$.

This occurs when the two lineages do not coalesce between the time of speciation, $t_s$, and introgression, $t_m$. Defining $t = t_m - t_s$, the probability of no coalescence is $p_{S_1} = e^{-t}$.

$S_2$ : The lineage configuration in the recipient branch at time $t_m$ consists of a single lineage, $\{AB\}$.

This occurs when the two lineages do coalesce between speciation, $t_s$, and introgression $t_m$. This happens with probability $p_{S_2} = 1 - e^{-t}$.

#### 2.2 Parent trees

The rate of introgression, $\gamma$, is defined as the probability that a lineage (branch) introgresses, and applies to each lineage that exists at the time of introgression independently. This means that we have to consider all possible scenarios of introgression within all possible lineage configurations, as these configurations consist of different numbers of branches that do or do not introgress. These scenarios range from ones where none of the lineages introgress, to ones where all do, but also all the intermediate combinations. If the recipient branch subtends more than two species, we would also have to track the exact sub-topologies of each lineage configuration (see Section 3). These scenarios define all the parent-tree histories and their probabilities, which can then be used to compute the expected frequencies of gene tree topologies under the species network. For the example with only two species descending from the recipient branch, the possible parent trees are (labeled as in Figure S8):

1. There are two cases in which no lineages introgress, and therefore two different parent trees with no introgression (parent trees 1 and 2 in Figure S8). Each must be weighted differently, and neither exactly matches the speciation history implied by the species network (Figure S8A). These are divided according to the lineage configuration at time $t_m$.

   When the lineage configuration is $S_1 = (A, B)$, the probability that no lineages introgress is $(1-\gamma)^2$, given that there are two independent lineages that could introgress. We represent the constraint of having no coalescence between $A$ and $B$ before $t_m$ in this configuration by having their time of divergence in parent tree 1 be $t_m$ (Figure S8B).

   Combining this with the probability of $S_1$, the probability of parent tree 1 is $p_1 = (1-\gamma)^2 e^{-t}$.

2. If there is no introgression and the lineage configuration is $S_2 = AB$, the probability that no lineages introgress is $(1-\gamma)$, given that there is just one possible lineage. We represent the constraint of only one lineage in this configuration as a single branch leading to tip $AB$ in parent tree 2 (Figure S8C). If needed, the coalescence time between $A$ and $B$, conditional on this occurring by $t_m$, can be calculated using equation A.7 in (Mendes and Hahn 2018).

   Combining this with the probability of $S_2$, the probability of parent tree 2 is $p_2 = (1-\gamma)(1-e^{-t})$.

3. For the case in which only lineage $A$ introgresses, the lineage configuration must be $S_1$. The probability that exactly one of two lineages introgesses in this configuration is $\gamma(1-\gamma)$.

   Combining this with the probability of $S_1$, the probability of parent tree 3 is $p_3 = \gamma(1-\gamma)e^{-t}$.

4. For the case in which only lineage $B$ introgresses, the lineage configuration must be $S_1$. By symmetry, this parent tree has the same probability as for parent tree 3, so the probability of parent tree 4 is $p_4 = \gamma(1-\gamma)e^{-t}$.

5. For the case in which $A$ and $B$ have already coalesced and therefore introgress together, the lineage configuration must be $S_2$. Since there is just one lineage at time $t_m$ in this configuration, the probability that it introgresses is simply $\gamma$. Again, this is represented as a single branch in parent tree 5 (Figure S8F).

   Combining this with the probability of $S_2$, the probability of parent tree 5 is $p_5 = \gamma(1-e^{-t})$.

6. Finally, there can be a scenario in which lineages $A$ and $B$ at the same locus introgress separately, for which the lineage configuration must be $S_1$. The probability of the two lineages introgressing is $\gamma^2$. Because they introgress at the same time and can coalesce with donor lineages independently, parent tree 6 is represented as a polytomy at time $t_m$ (Figure S8G).

   Combining this with the probability of $S_1$, the probability of parent tree 6 is $p_6 = \gamma^2 e^{-t}$.

#### 2.3 Expected gene tree frequencies

Summarizing the probabilities (i.e. weights) of the different parent trees, we have:

- $p_1 = (1-\gamma)^2 e^{-t}$,
- $p_2 = (1-\gamma)(1-e^{-t})$,
- $p_3 = \gamma(1-\gamma)e^{-t}$,
- $p_4 = \gamma(1-\gamma)e^{-t}$,
- $p_5 = \gamma(1-e^{-t})$,
- $p_6 = \gamma^2 e^{-t}$.

Given the six parent trees and weights, the expected frequency of any given topology $\tau$ under this species network can be computed as

$$P(\tau \mid \text{Species network}) = p_1 P(\tau|1) + p_2 P(\tau|2) + p_3 P(\tau|3) + p_4 P(\tau|4) + p_5 P(\tau|5), + p_6 P(\tau|6),$$

where $P(\tau|k)$ is the probability of a gene tree given parent tree $k$ under the multispecies coalescent. More generally,

$$P(\tau \mid \text{Species network}) = \sum_{k \in \{K\}} p_k P(\tau|k),$$

for the case where there are $K$ different parent trees. An example calculation using the expected frequency of discordant attachments is shown in Figure S9.

### 3 Generalizing to introgression with more descendants

Although none of the calculations in the main text are for trees with introgression into a branch with more than two descendants, here we illustrate the general logic of all such models. To compute the expected frequency of a topology under the species network when there is introgression into an internal branch, we proceed in three steps:

1. Identify all possible lineage configurations present in the recipient branch at the introgression time.
2. For each configuration, enumerate all possible subsets of lineages that can introgress and compute their probabilities.
3. Compute the expected topology frequencies under each resulting parent-tree history, and combine them as a weighted average using the probabilities of the corresponding scenarios.

#### 3.1 Lineage Configurations

We first revisit the example where there are two species below the recipient branch. In this case there are only two possible lineage configurations at time $t_m$:

$S_1$ : The configuration consists of two lineages, $\{A, B\}$.

$S_2$ : The configuration consists of a single lineage, $\{AB\}$.

Thus, in the two-descendant case, the number of lineages completely determines the lineage configuration. This is no longer true when the recipient branch has three or more descendant taxa, because different coalescent events can lead to different configurations with the same number of lineages.

To illustrate this, let us consider a recipient branch whose descendant subtree has topology $((A, B), C)$. In this case, the lineage configuration at time $t_m$ may consist of one, two, or three lineages, depending on how many coalescent events have occurred. However, it is also necessary to distinguish which coalescent events have occurred. Taking this into account, the seven possible configurations are the following, labeled as in Figure A1:

$S_1$ : The configuration consists of three separate lineages, $\{A, B, C\}$.

$S_2$ : The configuration consists of two lineages, $\{AB, C\}$.

$S_3$ : The configuration consists of two lineages, $\{A, BC\}$.

$S_4$ : The configuration consists of two lineages, $\{AC, B\}$.

$S_5$ : The configuration consists of a single lineage with internal structure $((A, B), C)$, namely $\{((A, B), C)\}$.

$S_6$ : The configuration consists of a single lineage with internal structure $(A, (B, C))$, namely $\{(A, (B, C))\}$.

$S_7$ : The configuration consists of a single lineage with internal structure $((A, C), B)$, namely $\{((A, C), B)\}$.

We now generalize this counting problem to a recipient branch with $N$ descendant taxa, so that we can see how many lineage configurations there are in any scenario.

Let $\mathcal{L}_N$ denote the set of possible lineage configurations at time $t_m$ when the recipient branch has $N$ descendant taxa and $L_N$ be the number of elements in this set (i.e $L_N = |\mathcal{L}_N|$). From the examples above, we have

$$L_2 = 2 \qquad \text{and} \qquad L_3 = 7.$$

Moreover, it is immediately true that

$$L_1 = 1.$$

Similarly, let us define $\mathcal{L}_{N,k}$ as the subset of $\mathcal{L}_N$ consisting of configurations with exactly $k$ lineages at time $t_m$, and let

$$L_{N,k} = |\mathcal{L}_{N,k}|.$$

Thus,

$$\mathcal{L}_N = \bigsqcup_{k=1}^{N} \mathcal{L}_{N,k}, \quad L_N = \sum_{k=1}^{N} L_{N,k}.$$

For the two-descendant case,

$$L_{2,1} = 1, \qquad L_{2,2} = 1,$$

so that

$$L_2 = L_{2,1} + L_{2,2} = 1 + 1 = 2.$$

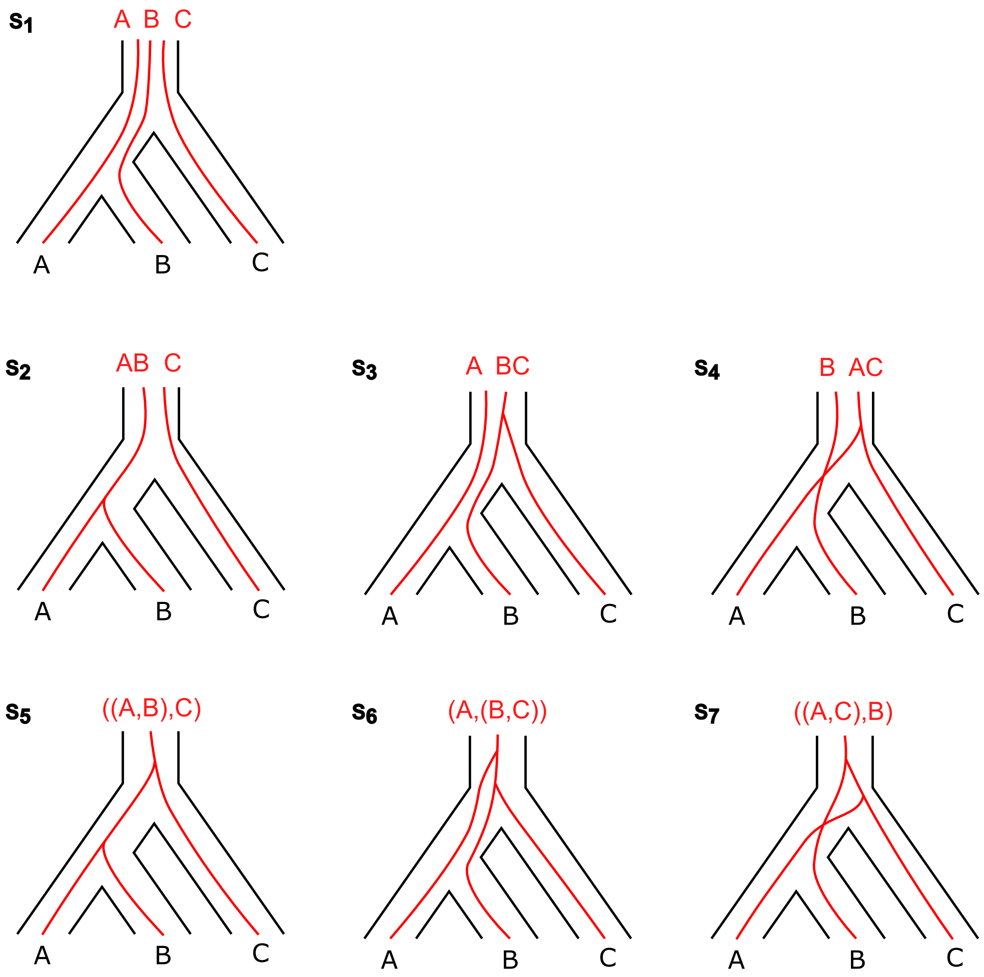


Figure A1: Possible lineage configurations for the recipient branch ((A,B),C).

For the three-descendant case,

$$L_{3,1} = 3, \qquad L_{3,2} = 3, \qquad L_{3,3} = 1,$$

and therefore

$$L_3 = L_{3,1} + L_{3,2} + L_{3,3} = 3 + 3 + 1 = 7.$$

We also adopt the convention that

$$L_{N,k} = 0 \qquad \text{for all } k \notin \{1, \dots, N\}.$$

Now, let us prove that, for $N \geq 3$, the number of lineage configurations satisfies the recurrence

$$L_N = (2N - 3)L_{N-1} + L_{N-2},$$

with initial conditions $L_1 = 1$ and $L_2 = 2$.

*Proof.* We partition the set of lineage configurations for $N$ taxa with exactly $k$ lineages in two classes, according to the position of the $N$th taxon:

$\mathcal{A}_{N,k}$: configurations in which the $N$th taxon forms a lineage by itself;

$\mathcal{B}_{N,k}$: configurations in which the $N$th taxon belongs to a lineage containing at least one other taxon.

Thus,

$$\mathcal{L}_{N,k} = \mathcal{A}_{N,k} \sqcup \mathcal{B}_{N,k},$$

$$L_{N,k} = |\mathcal{A}_{N,k}| + |\mathcal{B}_{N,k}|.$$

There is a natural bijection between $\mathcal{A}_{N,k}$ and the set of lineage configurations for $N - 1$ taxa with $k - 1$ lineages at $t_m$. Indeed, deleting the singleton lineage containing the $N$th taxon, one goes from an element in $\mathcal{A}_{N,k}$ to a lineage configuration for $N-1$ taxa with $k-1$ lineages. Conversely, any configuration given any member of the latter set can be extended to a configuration in $\mathcal{A}_{N,k}$ by adding the $N$th taxon as a new lineage. Therefore,

$$|\mathcal{A}_{N,k}| = L_{N-1,k-1}.$$

It remains to count the number of configurations in $\mathcal{B}_{N,k}$. To construct such a configuration, we start with a lineage configuration of the first $N-1$ taxa with exactly $k$ lineages, and then attach taxon $N$ to one of the existing lineages. Considering a configuration of $N-1$ taxa with $k$ lineages, let $n_i$ be the number of taxa contained in the $i$th lineage, for $i = 1, \dots, k$. Since the configuration contains $N-1$ taxa, we have

$$\sum_{i=1}^{k} n_i = N - 1.$$

For a lineage whose internal structure is a rooted binary tree with $M$ taxa, there are $2M-1$ possible ways to attach a new taxon. So the possible ways to attach the $N$th taxa to one of the existing lineages is

$$\sum_{i=1}^{k}(2n_i - 1) = 2\sum_{i=1}^{k} n_i - k = 2(N-1) - k = 2N - k - 2.$$

Thus, each lineage configuration of the first $N-1$ taxa with $k$ lineages gives rise to $2N-k-2$ configurations in $\mathcal{B}_{N,k}$. Therefore,

$$|\mathcal{B}_{N,k}| = (2N-k-2)L_{N-1,k}.$$

Combining the counts for the two classes, we obtain

$$L_{N,k} = L_{N-1,k-1} + L_{N-1,k}(2N-k-2).$$

Now, we sum over $k$ to get $L_N$

$$L_N = \sum_{k=1}^{N} L_{N,k} = \sum_{k=1}^{N} L_{N-1,k-1} + \sum_{k=1}^{N}(2N-k-2)L_{N-1,k}$$

$$L_N = L_{N-1} + 2NL_{N-1} - 2L_{N-1} - \sum_{k=1}^{N} kL_{N-1}$$

$$L_N = (2N-1)L_{N-1} - \sum_{k=1}^{N} kL_{N-1,k}$$

The only thing remaining is to simplify $\sum_{k=1}^{N} kL_{N-1,k}$.

More generally, the number $\sum_{k=1}^{m} kL_{m,k}$ counts the number of lineage configurations of $m$ taxa with one of these lineages marked, since each lineage configuration is counted as many times as lineages it has. Let us now consider the set of these configurations in which the marked lineage does not contain the $m$th taxon and call it $\mathcal{M}_m$. Using the same reasoning, we get

$$|\mathcal{M}_m| = \sum_{k=1}^{m}(k-1)L_{m,k}.$$

We claim that there is a bijection between $\mathcal{M}_m$ and the set of lineage configurations of $m$ taxa in which the $m$th taxon does not form a singleton lineage. To see this, take a configuration of the latter set, since the $m$th taxon is not alone it has a sister subtree inside the lineage, consider this sister subtree and separate it from the rest of the lineage and mark the newly created lineage. As the marked lineage does not contain the $m$th taxon, this gives us an element of $\mathcal{M}_m$.

Conversely, given an element of $\mathcal{M}_m$, consider the marked lineage and the lineage that contains the $m$th taxon. Attach the first lineage to the second one, in such a way that the marked lineage becomes the sister subtree of the $m$th taxon. These operations are inverse to each other, so the claimed bijection holds.

The number of lineage configurations of $m$ taxa in which taxon $m$ forms a singleton lineage is $L_{m-1}$, since deleting that singleton lineage leaves an arbitrary configuration of the remaining $m-1$ taxa. Therefore, the number of configurations in which taxon $m$ is not alone is

$$L_m - L_{m-1}.$$

By the bijection above, we obtain

$$|\mathcal{M}_m| = L_m - L_{m-1}.$$

Hence,

$$\sum_{k=1}^{m} kL_{m,k} = \sum_{k=1}^{m}(k-1)L_{m,k} + L_m = L_m - L_{m-1} + L_m = 2L_m - L_{m-1}.$$

Finally, applying this identity with $m = N - 1$ and substituting it into the expression for $L_N$ we get

$$L_N = (2N-1)L_{N-1} - 2L_{N-1} + L_{N-2} = (2N-3)L_{N-1} + L_{N-2},$$

which is the recurrence relation needed to calculate the number of lineage configurations for any $N$. □

#### 3.2 Parent trees

Once the possible lineage configurations at time $t_m$ have been identified, they can be used to describe the corresponding parent trees. Returning to the two-descendant example, we have already seen that it gives rise to six different parent trees (Figure S8).

We now work out the three-descendant case. From the previous section, the three-descendant case has lineage configurations distributed as follows:

$$L_{3,1} = 3, \qquad L_{3,2} = 3, \qquad L_{3,3} = 1.$$

Each lineage configuration gives rise to a different number of parent trees, depending on the number of lineages present at time $t_m$. For the three-descendant case, this gives:

$\mathcal{L}_{3,1}$: Each configuration in this set has one lineage. Therefore, there are two possible scenarios: either the lineage introgresses or it does not. Hence, each configuration in this set gives rise to 2 parent trees.

$\mathcal{L}_{3,2}$: Each configuration in this set has two lineages. Therefore, there are four possible scenarios:

- neither lineage introgresses;
- the first lineage introgresses and the second one does not;
- the first lineage does not introgress and the second one does;
- both lineages introgress.

Hence, each configuration in this set gives rise to 4 parent trees.

$\mathcal{L}_{3,3}$: Each configuration in this set has three lineages. Therefore, there are 8 possible scenarios, one for each possible subset of lineages introgressing. Hence, each configuration in this set gives rise to 8 parent trees.

Let $P_3$ denote the total number of parent trees for the three-descendant case. Then

$$P_3 = L_{3,1} \cdot 2 + L_{3,2} \cdot 4 + L_{3,3} \cdot 8 = 3 \cdot 2 + 3 \cdot 4 + 1 \cdot 8 = 26.$$

An important observation from this computation is that if a configuration has $k$ lineages, then each lineage may either introgress or not introgress, giving $2^k$ possible parent trees.

We can use this observation to generalize to the case of $N$ descendant taxa. Let $P_N$ denote the total number of possible parent trees for a recipient branch with $N$ descendants. Since there are $L_{N,k}$ lineage configurations with exactly $k$ lineages, and each of them gives rise to $2^k$ parent trees, we have

$$P_N = \sum_{k=1}^{N} 2^k L_{N,k}.$$

#### 3.3 General expression for gene tree frequencies

Given the construction above, we can now write a general expression for the expected frequency of a given topology, $\tau$, under a species tree with introgression. Let $\mathcal{L}_N$ be the set of possible lineage configurations at time $t_m$ when the recipient branch has $N$ descendant taxa. Then

$$P(\tau) = \sum_{L \in \mathcal{L}_N} P(L) \left[ \sum_{A \subseteq L} \gamma^{|A|}(1-\gamma)^{|L|-|A|}\, P(\tau \mid T(L, A)) \right]$$

Here,

$$
\begin{aligned}
N &= \text{number of taxa below the recipient branch,} \\
\mathcal{L}_N &= \text{set of lineage configurations at time } t_m, \\
L &= \text{a lineage configuration in } \mathcal{L}_N, \\
A &= \text{subset of lineages in the lineage configuration } L, \\
T(L, A) &= \text{parent tree obtained from } L \text{ when the lineages in } A \text{ introgress,} \\
\gamma &= \text{probability of introgression.}
\end{aligned}
$$

The outer summation has $|\mathcal{L}_N| = L_N$ terms, one for each possible lineage configuration. For a fixed lineage configuration $L$, the inner summation ranges over all subsets of the lineages in $L$. Thus, if $L$ contains $|L|$ lineages, the inner summation has $2^{|L|}$ terms. Consequently, the total number of terms in the double summation is

$$\sum_{L \in \mathcal{L}_N} 2^{|L|}.$$

This is precisely the number of possible parent trees, which we denoted by $P_N$. Therefore,

$$P_N = \sum_{L \in \mathcal{L}_N} 2^{|L|} = \sum_{k=1}^{N} 2^k L_{N,k}.$$

Table A1 shows the values of $L_N$ and $P_N$ for $N = 1, \ldots, 4$. These values illustrate the rapid growth of the number of terms in the general expression. When $N = 4$, there are already $P_4 = 154$ parent trees. This rapid increase makes direct computations under the general model computationally infeasible as $N$ grows large.

| **Number of taxa** ($N$) | **Lineage configurations** ($L_N$) | **Parent trees** ($P_N$) |
|---|---|---|
| 1 | 1 | 2 |
| 2 | 2 | 6 |
| 3 | 7 | 26 |
| 4 | 37 | 154 |

Table A1: Number of lineage configurations $L_N$ and parent trees $P_N$ for $N = 1, \ldots, 4$.
