## Supplementary material for "New approaches to detecting and characterizing introgression in large species trees": S3_Table.docx

**Table S3**. Probability of each gene tree topology for species tree (((*Z,Y*),*X*),*W*) under the multispecies coalescent (from Table V and Equation 2 in Rosenberg [2002]).

Both the speciation history (parent tree 1) and the introgression history (parent tree 2) shown in Figure 3 of the main text have this same tree shape, though the length of the two internal branches differs between them. For either history, we denote the length (in coalescent units) of the more recent internal branch as *a* and the older internal branch as *b*. The length of *a* is therefore *t*_2_- *t*_1_ in parent tree 1 and *t*_1_- *t*_m_ in parent tree 2, while the length of *b* is *t*_3_- *t*_2_ in parent tree 1 and *t*_2_- *t*_1_ in parent tree 2.

| Topology | Probability |
| --- | --- |
| 1. ((*Z,Y*),(*X,W*)) | $\frac{1}{3}e^{-b}-\frac{1}{6}e^{-\left( a+b \right)}-\frac{1}{18}e^{-(3a+b)}$ |
| 2. ((*Z,X*),(*Y,W*)) | $\frac{1}{6}e^{-\left( a+b \right)}-\frac{1}{18}e^{-(3a+b)}$ |
| 3. ((*Y,X*),(*Z,W*)) | $\frac{1}{6}e^{-\left( a+b \right)}-\frac{1}{18}e^{-(3a+b)}$ |
| 4. (((*Z,Y*),*X*),*W*) | $1-\frac{2}{3}e^{-a}-\frac{2}{3}e^{-b}+\frac{1}{3}e^{-\left( a+b \right)}+\frac{1}{18}e^{-(3a+b)}$ |
| 5. (((*Z,Y*),*W*),*X*) | $\frac{1}{3}e^{-b}-\frac{1}{6}e^{-\left( a+b \right)}-\frac{1}{9}e^{-(3a+b)}$ |
| 6. (((*Z,X*),*Y*),*W*) | _­_$\frac{1}{3}e^{-a}-\frac{1}{3}e^{-\left( a+b \right)}+\frac{1}{18}e^{-(3a+b)}$ |
| 7. (((*Z,X*),*W*),*Y*) | $\frac{1}{6}e^{-(a+b)}-\frac{1}{9}e^{-(3a+b)}$ |
| 8. (((*Z,W*),*Y*),*X*) | $\frac{1}{18}e^{-(3a+b)}$ |
| 9. (((*Z,W*),*X*),*Y*) | $\frac{1}{18}e^{-(3a+b)}$ |
| 10. (((*Y,X*),*Z*),*W*) | _­_$\frac{1}{3}e^{-a}-\frac{1}{3}e^{-\left( a+b \right)}+\frac{1}{18}e^{-(3a+b)}$ |
| 11. (((*Y,X*),*W*),*Z*) | $\frac{1}{6}e^{-(a+b)}-\frac{1}{9}e^{-(3a+b)}$ |
| 12. (((*Y,W*),*Z*),*X*) | $\frac{1}{18}e^{-(3a+b)}$ |
| 13. (((*Y,W*),*X*),*Z*) | $\frac{1}{18}e^{-(3a+b)}$ |
| 14. (((*X,W*),*Z*),*Y*) | $\frac{1}{18}e^{-(3a+b)}$ |
| 15. (((*X,W*),*Y*),*Z*) | $\frac{1}{18}e^{-(3a+b)}$ |
