## Supplementary figures and images for "New approaches to detecting and characterizing introgression in large species trees"

### S1_Fig.tiff

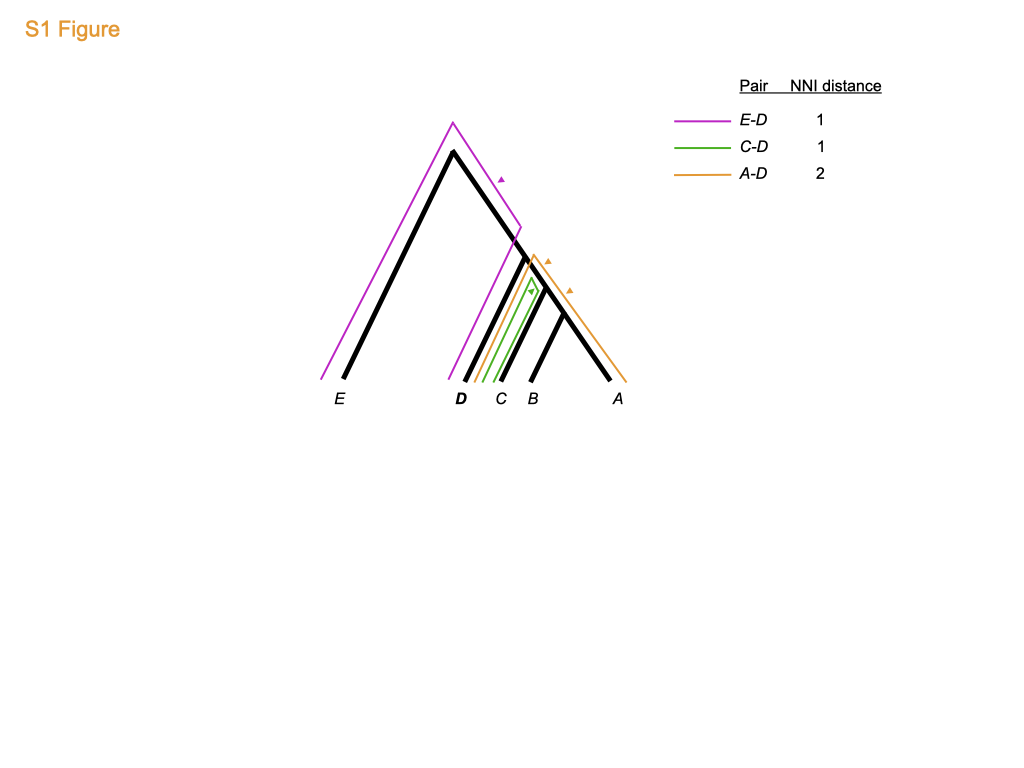

### S1_Table.tiff

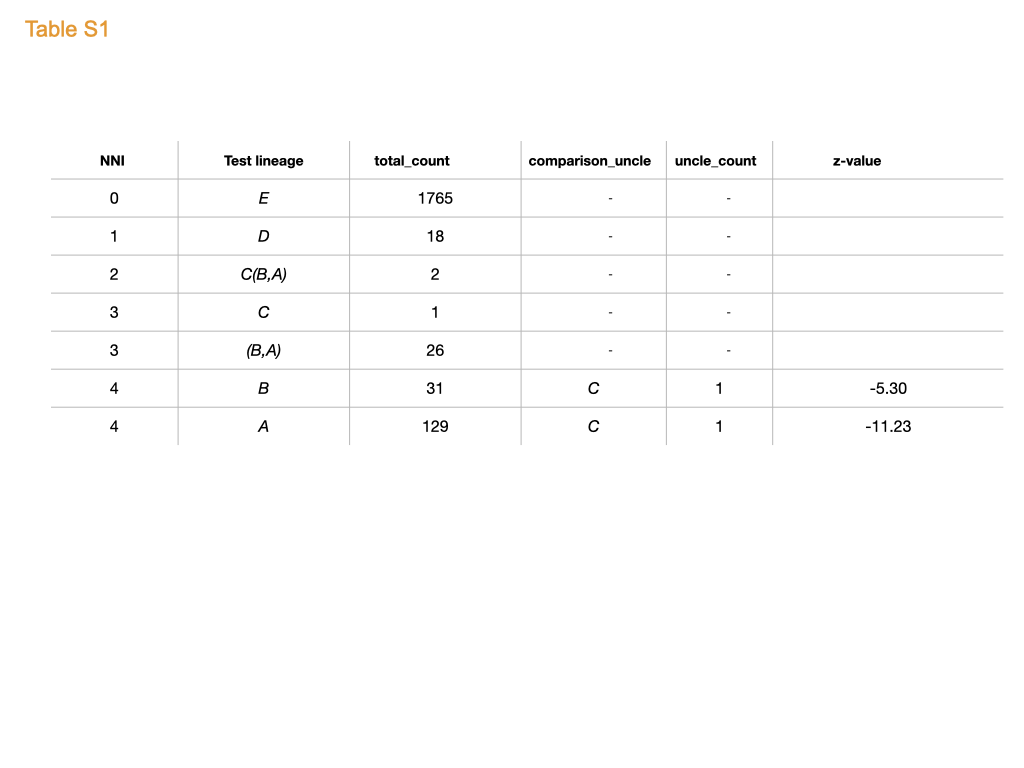

### S2_Fig.tiff

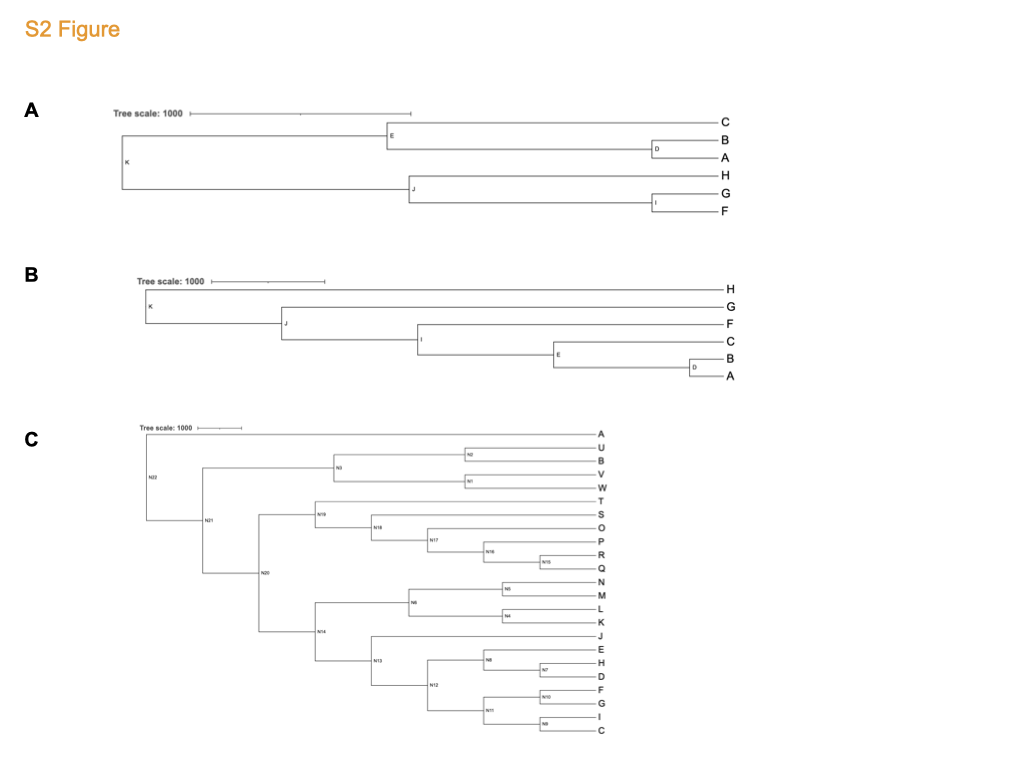

### S2_Table.tiff

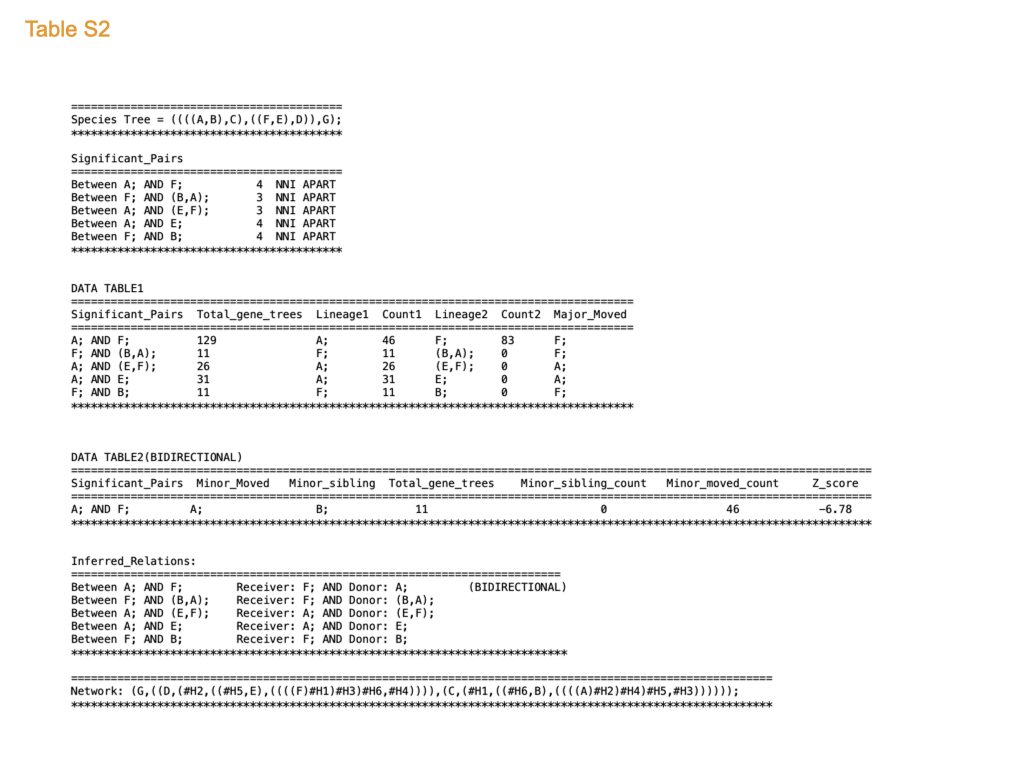

### S3_Fig.tiff

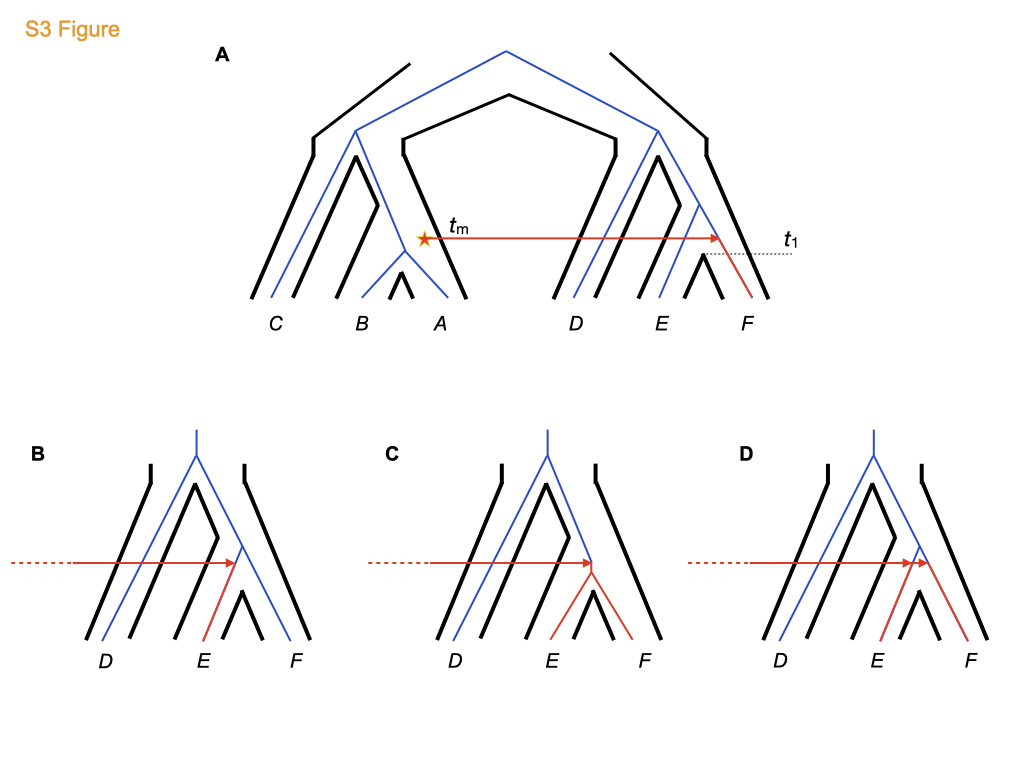

### S4_Fig.tiff

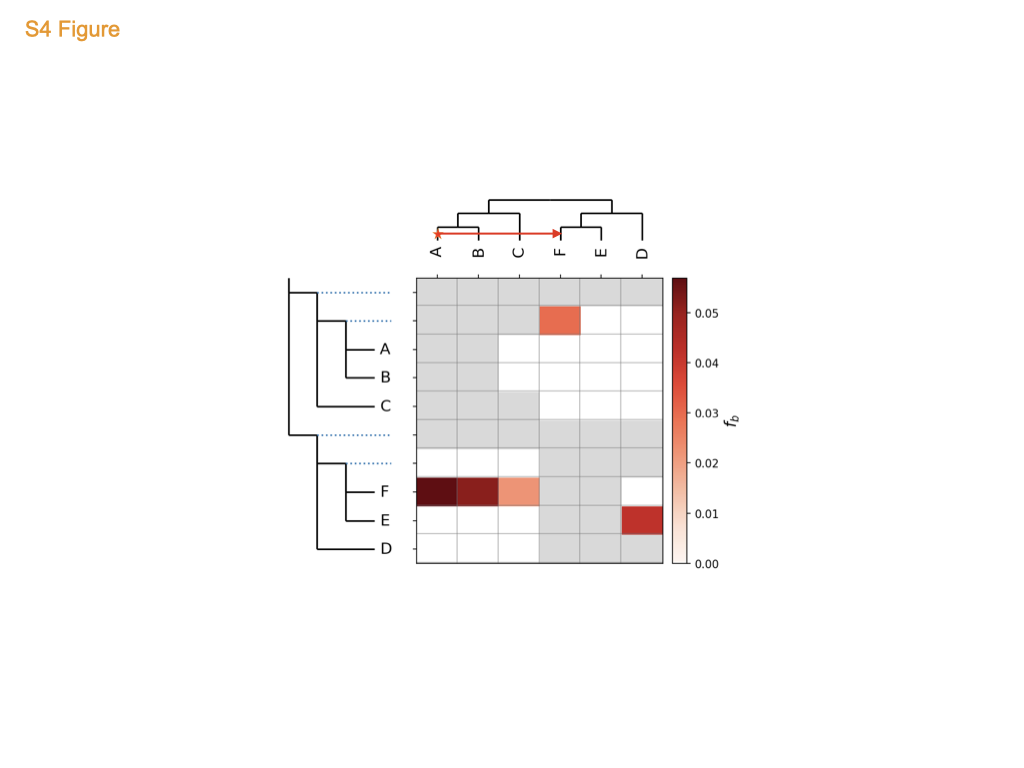

### S5_Fig.tiff

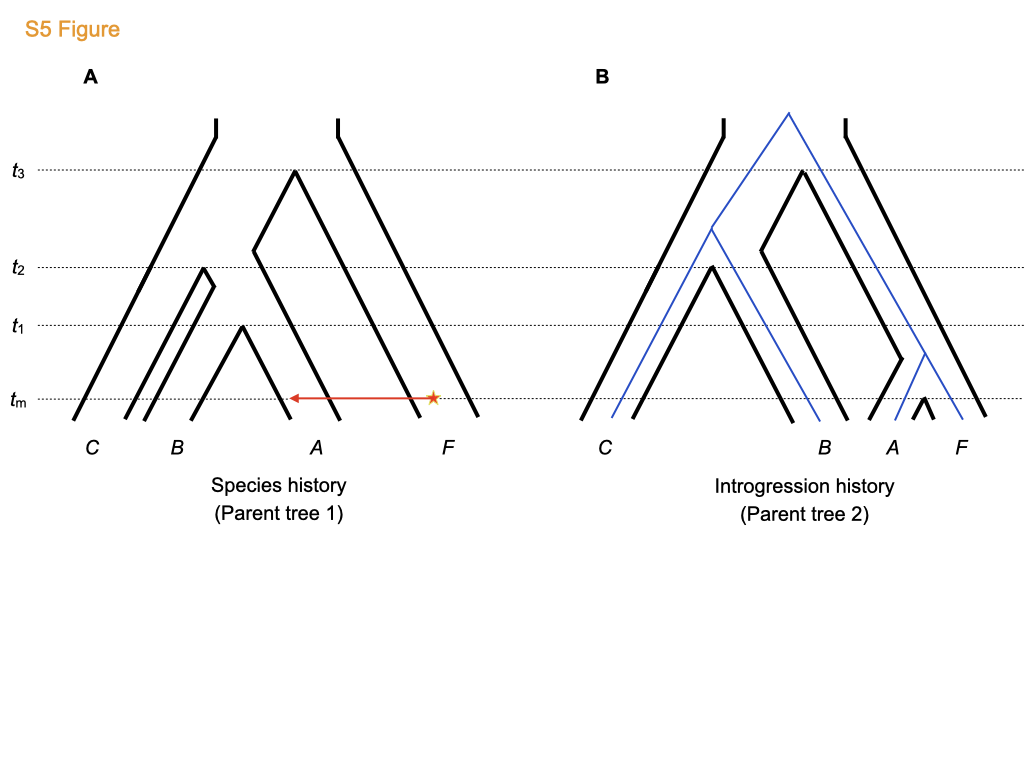

### S5_FigC.tiff

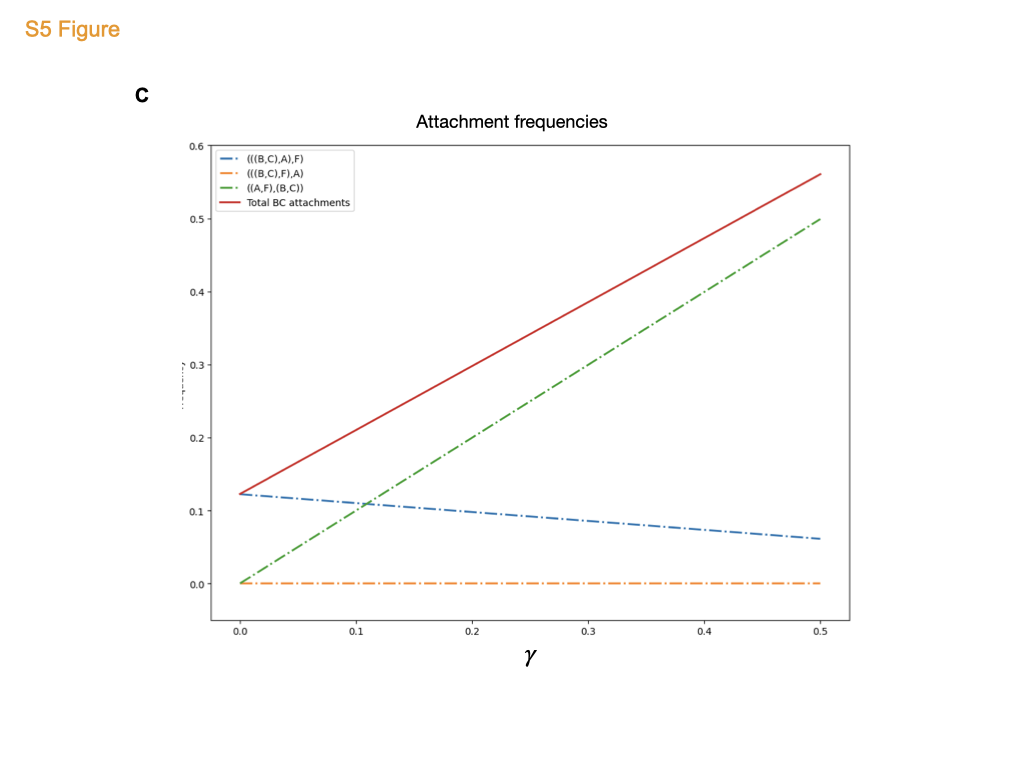

### S6_Fig.tiff

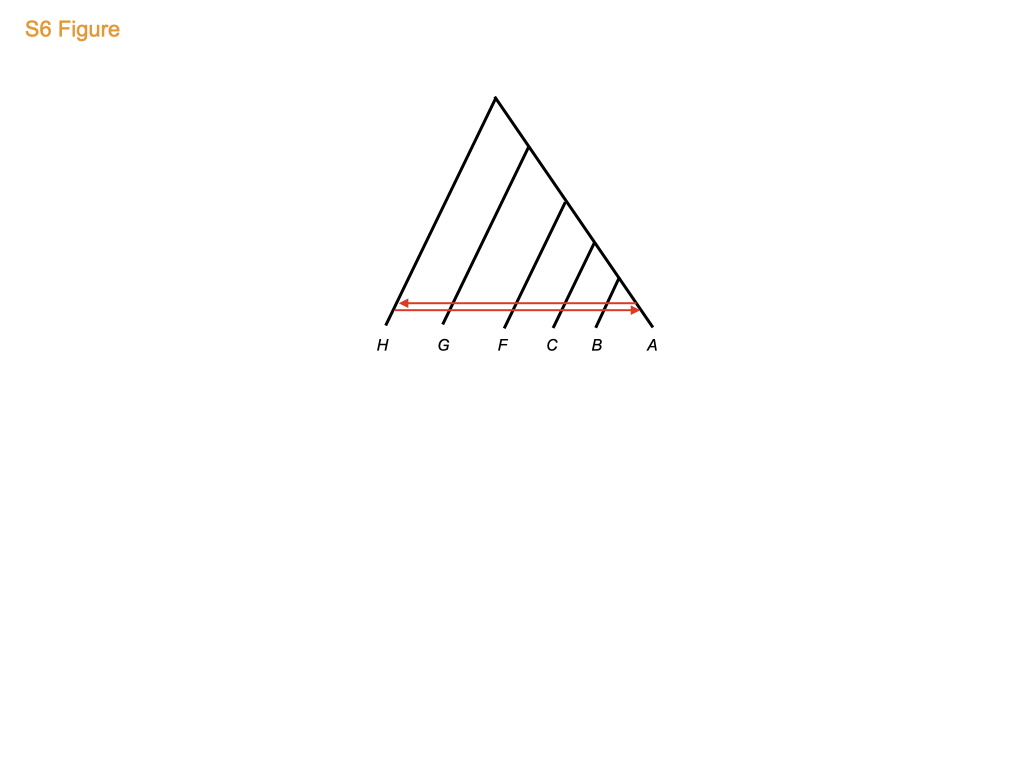

### S7_Fig.tiff

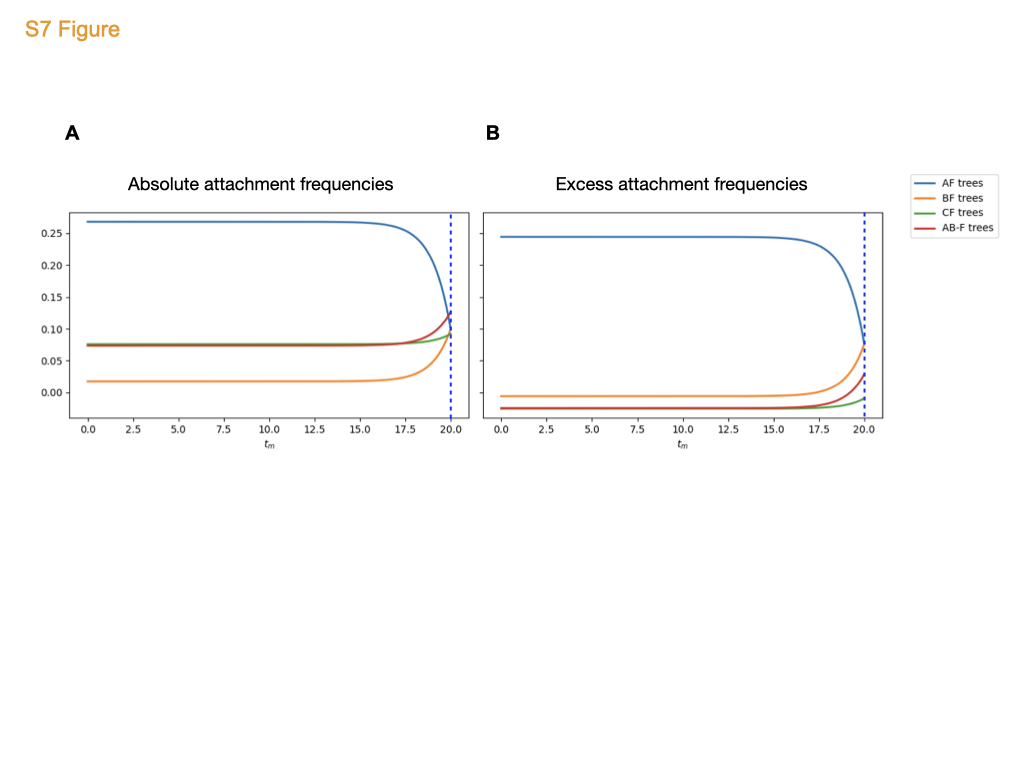

### S8_Fig.tiff

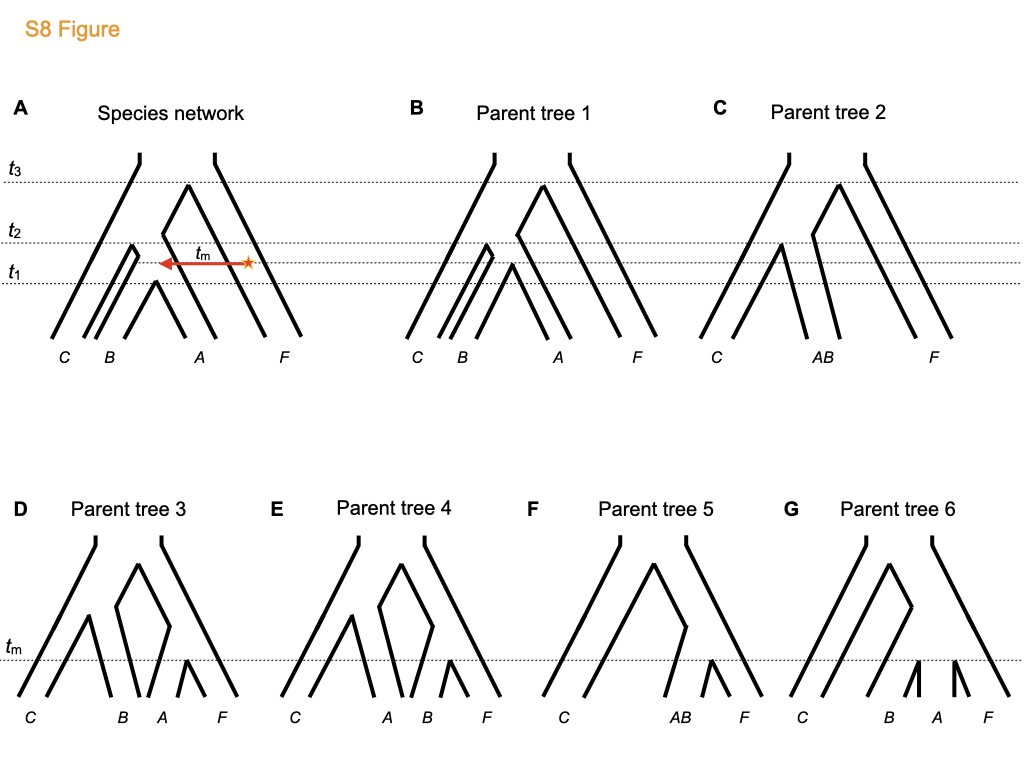

### S9_Fig.tiff

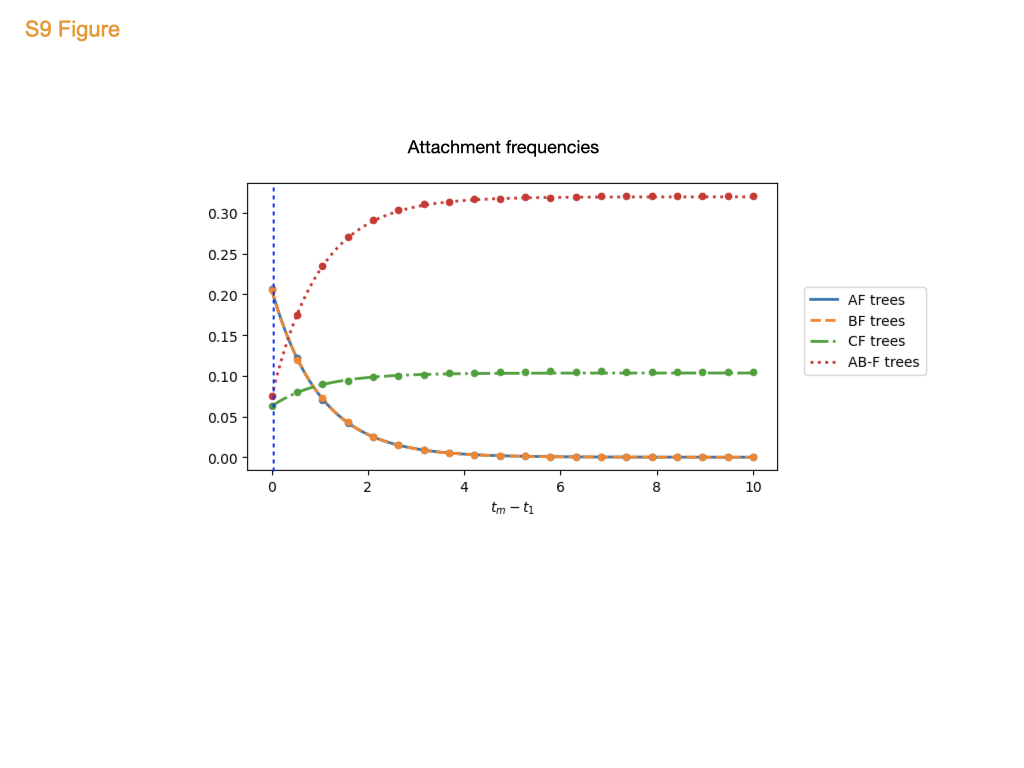

### S10_Fig.tiff

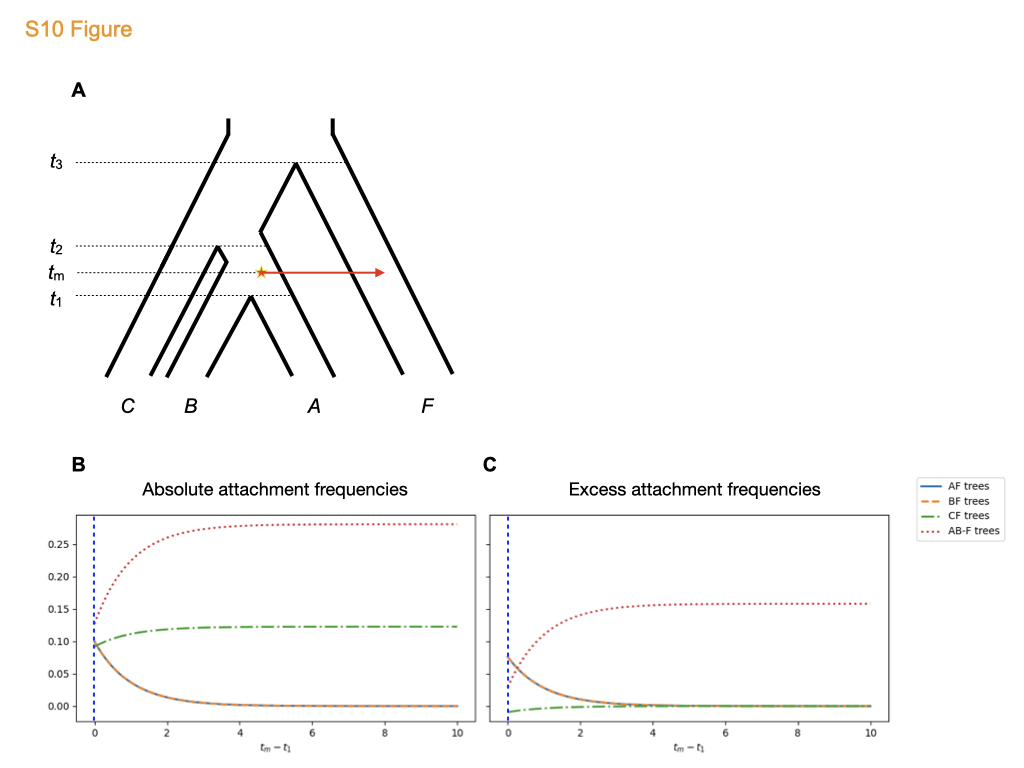
